# FMRIPrep: a robust preprocessing pipeline for functional MRI

**DOI:** 10.1101/306951

**Authors:** Oscar Esteban, Christopher J. Markiewicz, Ross W. Blair, Craig A. Moodie, A. Ilkay Isik, Asier Erramuzpe, James D. Kent, Mathias Goncalves, Elizabeth DuPre, Madeleine Snyder, Hiroyuki Oya, Satrajit S. Ghosh, Jessey Wright, Joke Durnez, Russell A. Poldrack, Krzysztof J. Gorgolewski

**Author notes:** **For correspondence:** (OE); (KG). Contributed equally to this work (Russell A. Poldrack, Krzysztof J. Gorgolewski).

## Abstract

Preprocessing of functional MRI (fMRI) involves numerous steps to clean and standardize data before statistical analysis. Generally, researchers create *ad hoc* preprocessing workflows for each new dataset, building upon a large inventory of tools available for each step. The complexity of these workflows has snowballed with rapid advances in MR data acquisition and image processing techniques. We introduce *fMRIPrep*, an analysis-agnostic tool that addresses the challenge of robust and reproducible preprocessing for task-based and resting fMRI data. *FMRIPrep* automatically adapts a best-in-breed workflow to the idiosyncrasies of virtually any dataset, ensuring high-quality preprocessing with no manual intervention. By introducing visual assessment checkpoints into an iterative integration framework for software-testing, we show that *fMRIPrep* robustly produces high-quality results on a diverse fMRI data collection comprising participants from 54 different studies in the OpenfMRI repository. We review the distinctive features of *fMRIPrep* in a qualitative comparison to other preprocessing workflows. We demonstrate that *fMRIPrep* achieves higher spatial accuracy as it introduces less uncontrolled spatial smoothness than commonly used preprocessing tools. *FMRIPrep* has the potential to transform fMRI research by equipping neuroscientists with a high-quality, robust, easy-to-use and transparent preprocessing workflow which can help ensure the validity of inference and the interpretability of their results.

---

Functional magnetic resonance imaging (fMRI) is a commonly used technique to map human brain activity^1^. However, the blood-oxygen-level dependent (BOLD) signal measured by fMRI is typically mixed with many non-neural sources of variability^2^. Preprocessing identifies the nuisance sources and reduces their effect on the data^3^. Other major preprocessing steps^4^ deal with particular imaging artifacts and the anatomical location of signals. For instance, slice-timing^5^ correction (STC), head-motion correction (HMC), and susceptibility distortion correction (SDC) address particular artifacts; while co-registration, and spatial normalization are concerned with signal location (see Online Methods, sec. Preprocessing of fMRI in a nutshell, for a summary). Extracting a signal that is most faithful to the underlying neural activity is crucial to ensure the validity of inference and interpretability of results^6^. Faulty preprocessing may lead to the interpretation of noise patterns as signals of interest. For example, Power et al. demonstrated that unaccounted-for head-motion can generate spurious and systematic correlations in resting-state fMRI^7^, which would be interpreted as functional connectivity. An illustration of failed spatial normalization familiar to most researchers is finding significant activation outside of the brain. Other preprocessing choices may result in the removal of signal originating from brain activity. The ongoing debate on the need for regressing out global signals^2,8,9^ reflects just such concerns. Thus, a primary goal of preprocessing is to reduce sources of Type I errors without inducing excessive Type II errors.

Workflows for preprocessing fMRI produce two broad classes of outputs: *preprocessed* data (as opposed to raw, original data) and measurements of experimental *confounds* for use in later modeling. Preprocessed data generally include new fMRI time-series after the application of retrospective signal correction and filtering algorithms. In addition, these data are typically resampled onto a target space appropriate for analysis, such as a standardized anatomical reference. The *confounds* are additional time-series such as physiological recordings and estimated noise sources that are useful for analysis (e.g. they can be applied as nuisance regressors). Some commonly used confounds include: motion parameters, framewise displacement (FD^7^), spatial standard deviation of the data after temporal differencing (DVARS^7^), global signals, etc. Preprocessing may include further steps for denoising and estimation of confounds. For instance, dimensionality reduction methods based on principal components analysis (PCA) or independent components analysis (ICA), such as component-based noise correction *(CompCor*^10^) or automatic removal of motion artifacts (ICA-AROMA^11^).

The neuroimaging community is well equipped with tools that implement the majority of the individual steps of preprocessing described so far. These tools are readily available within software packages including AFNI^12^, ANTs^13^, FreeSurfer^14^, FSL^15^, Nilearn^16^, or SPM^17^. Despite the wealth of accessible software and multiple attempts to outline best practices for preprocessing^2,4,6,18^, the large variety of data acquisition protocols have led to the use of *ad hoc* pipelines customized for nearly every study; for example, Carp^19^ found 223 unique analysis workflows across 241 fMRI studies. Thus, current preprocessing workflows offer a poor trade-off between the quality of results and robust, consistent performance on datasets other than those that they were built for. Alternatively, researchers can adopt the acquisition protocols defined by large neuroimaging consortia like the Human Connectome Project (HCP^20^) or the UK Biobank^21^, which then allows the use of their preprocessing pipelines^22,23^ developed for those studies. Since these pipelines are optimized for particular data acquisition protocols, they are not applicable to datasets acquired using different protocols. In practice, the neuroimaging community lacks a preprocessing workflow that reliably provides high-quality and consistent results on arbitrary datasets.

Here we introduce *fMRIPrep*, a preprocessing workflow for task-based and resting-state fMRI. *FMRIPrep* is built around four driving principles: 1) **robustness** to the idiosyncrasies of the input dataset; 2) **quality** of preprocessing outcomes; 3) **transparency** to encourage the scrutiny of preprocessing results for quality, and to facilitate accurate communication of the methods; and 4) **ease-of-use** with the minimization of manual intervention. *FMRIPrep* is robust by virtue of a flexible, self-adapting architecture that combines tools from existing neuroimaging analysis packages. Tools for each processing operation are selected through an evidence-driven and community-informed optimization process. Here we also report a comprehensive evaluation of the workflow on a large and heterogeneous subsample of the OpenfMRI repository, to quantify robustness and quality of the results. This evaluation leverages the comprehensive visual reports generated by *fMRIPrep*, which facilitate assessment and curation of the results. These reports contribute to the “glass-box” philosophy with which the software was developed; rather than hiding a complex set of operations within a monolithic black box, *fMRIPrep* exposes interim results at multiple steps to encourage active engagement by the scientist.

## RESULTS

*FMRIPrep* is a robust and convenient tool for researchers and clinicians to prepare both task-based and resting-state fMRI data for analysis. Its outputs enable a broad range of applications, including within-subject analysis using functional localizers, voxel-based analysis, surface-based analysis, task-based group analysis, resting-state connectivity analysis, and many others. In the following, we describe the overall architecture, software engineering principles, and a comprehensive validation of the tool.

**Figure 1.**
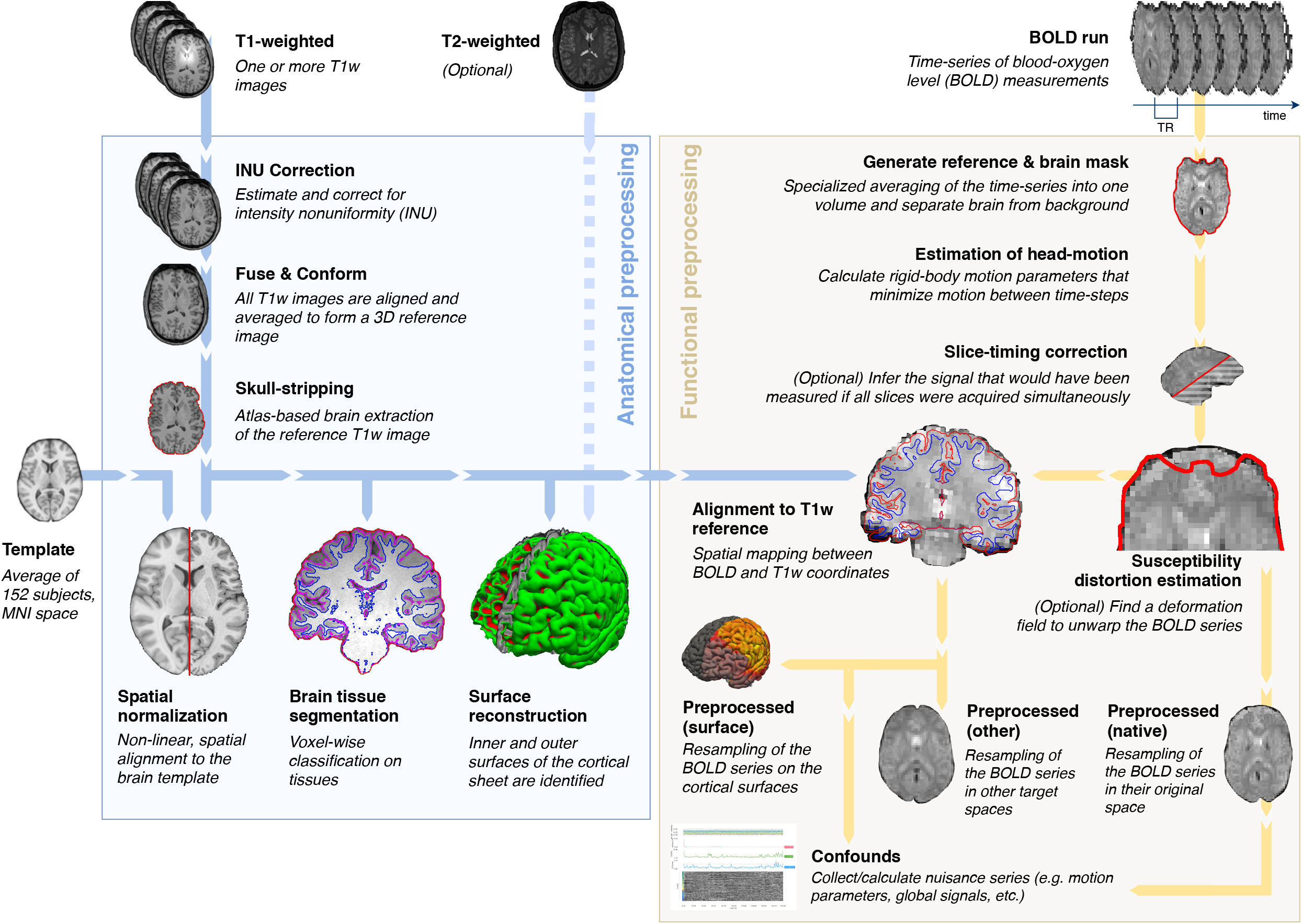
*FMRIPrep* is an fMRI preprocessing tool that adapts to the input dataset. Leveraging the Brain Imaging Data Structure (BIDS^24^), the software self-adjusts automatically, configuring the optimal workflow for the given input dataset. Thus, no manual intervention is required to locate the required inputs (one T1-weighted image and one BOLD series), read acquisition parameters (such as the repetition time –TR– and the slice acquisition-times) or find additional acquisitions intended for specific preprocessing steps (like field maps and other alternatives for the estimation of the susceptibility distortion). Outputs are easy to navigate due to compliance with the BIDS Extension Proposal for derived data (see Online Methods,*Figure S4*).

### A modular design allows for a flexible, adaptive workflow

The foundation of *fMRIPrep* is presented in Figure 1. The workflow is composed of sub-workflows that are dynamically assembled into different configurations depending on the input data. These building blocks combine tools from widely-used, open-source neuroimaging packages (see Table 1 for a summary). *Nipype*^25^ is used to stage the workflows and to deal with execution details (such as resource management). As presented in Figure 1, the workflow comprises two major blocks, separated into anatomical and functional MRI processing streams.

**Table 1.**
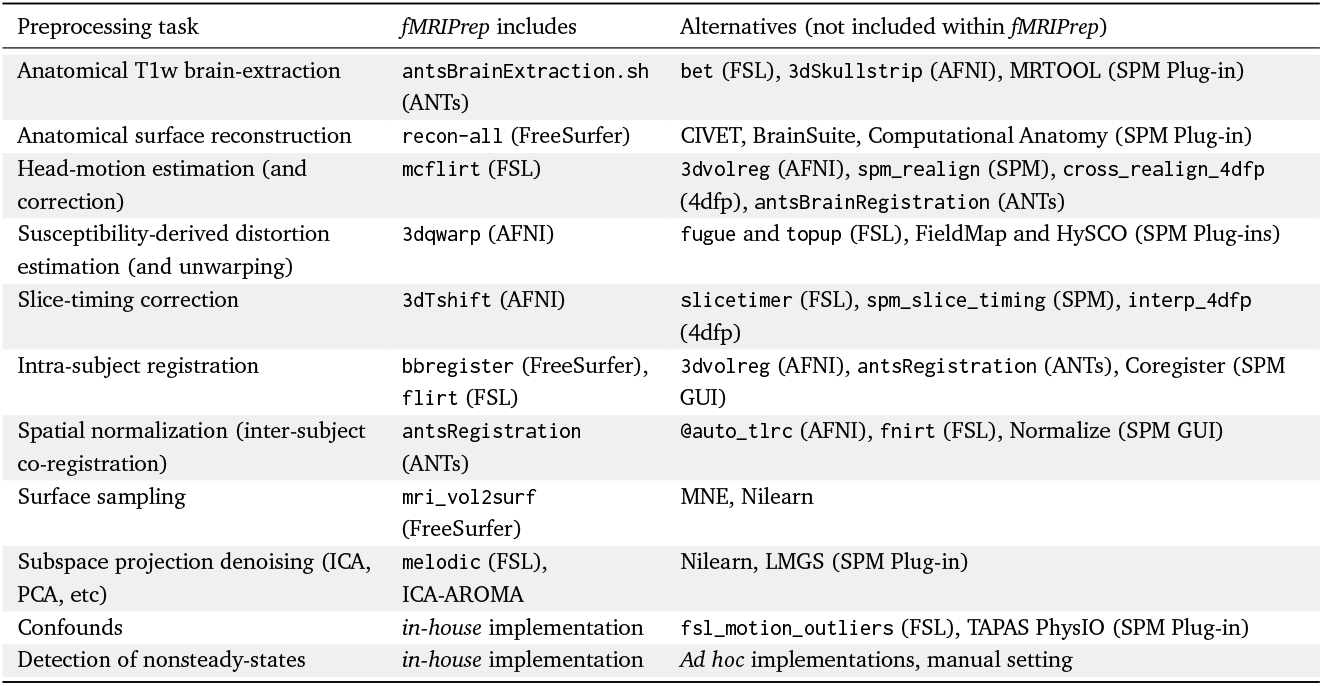
State-of-the-art neuroimaging offers a large catalog of readily available software tools. *FMRIPrep* integrates best-in-breed tools for each of the preprocessing tasks that its workflow covers, except for steps implemented as part of the development of *fMRIPrep (in-house* implementations). Tasks listed on the first column are described in detail in Online Methods, sec. Preprocessing of fMRI in a nutshell.

| Preprocessing task | *fMRIPrep* includes | Alternatives (not included within *fMRIPrep*) |
| --- | --- | --- |
| Anatomical T1w brain-extraction | antsBrainExtraction.sh (ANTs) | bet (FSL), 3dSkullstrip (AFNI), MRTOOL (SPM Plug-in) |
| Anatomical surface reconstruction | recon-all (FreeSurfer) | CIVET, BrainSuite, Computational Anatomy (SPM Plug-in) |
| Head-motion estimation (and correction) | mcflirt (FSL) | 3dvolreg (AFNI), spm_realign (SPM), cross_realign_4dfp (4dfp), antsBrainRegistration (ANTs) |
| Susceptibility-derived distortion estimation (and unwarping) | 3dqwarp (AFNI) | fugue and topup (FSL), FieldMap and HySCO (SPM Plug-ins) |
| Slice-timing correction | 3dTshift (AFNI) | slicetimer (FSL), spm_slice_timing (SPM), interp_4dfp (4dfp) |
| Intra-subject registration | bbregister (FreeSurfer), flirt (FSL) | 3dvolreg (AFNI), antsRegistration (ANTs), Coregister (SPM GUI) |
| Spatial normalization (inter-subject co-registration) | antsRegistration (ANTs) | @auto_tlrc (AFNI), fnirt (FSL), Normalize (SPM GUI) |
| Surface sampling | mri_vol2surf (FreeSurfer) | MNE, Nilearn |
| Subspace projection denoising (ICA, PCA, etc) | melodic (FSL), ICA-AROMA | Nilearn, LMGS (SPM Plug-in) |
| Confounds | *in-house* implementation | fsl_motion_outliers (FSL), TAPAS PhysIO (SPM Plug-in) |
| Detection of nonsteady-states | *in-house* implementation | *Ad hoc* implementations, manual setting |

#### Automatically understanding the input dataset

The Brain Imaging Data Structure (BIDS ^24^) allows *fMRIPrep* to precisely identify the structure of the input data and gather all the available metadata (e.g. imaging parameters). Online Methods, sec. The BIDS ecosystem describes how BIDS enables workflow flexibility. *FMRIPrep* reliably adapts to dataset irregularities such as missing acquisitions or runs through a set of heuristics. For instance, if only one participant of a sample lacks field-mapping acquisitions, *fMRIPrep* will by-pass the correction step for that one participant.

#### Preprocessing anatomical images

The T1-weighted (T1w) image is corrected for intensity nonuniformity (INU) using N4BiasFieldCorrection^26^ (ANTs), and skull-stripped using antsBrainExtraction.sh (ANTs). Skull-stripping is performed through coregistration to a template, with two options available: the OASIS template^27^ (default) or the NKI template^28^. Using visual inspection, we have found that this approach outperforms other common approaches, which is consistent with previous reports^22^. When several T1w volumes are found, the INU-corrected versions are first fused into a reference T1w map of the subject with mri_robust_template^29^ (FreeSurfer). Brain surfaces are reconstructed from the subject’s T1w reference (and T2-weighted images if available) using recon-all^30^ (FreeSurfer). The brain mask estimated previously is refined with a custom variation of a method (originally introduced in Mindboggle^31^) to reconcile ANTs-derived and FreeSurfer-derived segmentations of the cortical gray matter (GM). Both surface reconstruction and subsequent mask refinement are optional and can be disabled to save run time when surface-based analysis is not needed. Spatial normalization to the ICBM 152 Nonlinear Asymmetrical template^32^ (version 2009c) is performed through nonlinear registration with antsRegistration ^33^ (ANTs), using brain-extracted versions of both the T1w reference and the standard template. ANTs was selected due to its superior performance in terms of volumetric group level overlap^34^. Brain tissues –cerebrospinal fluid (CSF), white matter (WM) and GM– are segmented from the reference, brain-extracted T1w using fast^35^ (FSL).

#### Preprocessing functional runs

For every BOLD run found in the dataset, a reference volume and its skull-stripped version are generated using an in-house methodology (reported in Online Methods, sec. Particular processing elements of *fMRIPrep*). Then, head-motion parameters (volume-to-reference transform matrices, and corresponding rotation and translation parameters) are estimated using mcflirt^36^ (FSL). Among several alternatives (see Table 1), mcflirt is used because its results are comparable to other tools^37^ and it stores the estimated parameters in a format that facilitates the composition of spatial transforms to achieve one-step interpolation (see below). If slice timing information is available, BOLD runs are (optionally) slice time corrected using 3dTshift (AFNI^12^). When field map information is available, or the experimental “fieldmap-less” correction is requested (see Highlights of *fMRIPrep* within the neuroimaging context), SDC is performed using the appropriate methods (see Online Methods, *Figure S3*). This is followed by co-registration to the corresponding T1w reference using boundary-based registration^38^ with nine degrees of freedom (to minimize remaining distortions). If surface reconstruction is selected, *fMRIPrep* uses bbregister (FreeSurfer). Otherwise, the boundary based coregistration implemented in flirt (FSL) is applied. In our experience, bbregister yields the better results^38^ due to the high resolution and the topological correctness of the GM/WM surfaces driving registration. To support a large variety of output spaces for the results (e.g. the native space of BOLD runs, the corresponding T1w, FreeSurfer’s *fsaverage* spaces, the template used as target in the spatial normalization step, etc.), the transformations between spaces can be combined. For example, to generate preprocessed BOLD runs in template space (e.g. MNI), the following transforms are concatenated: head-motion parameters, the warping to reverse susceptibility-distortions (if calculated), BOLD-to-T1w, and T1w-to-template mappings. The BOLD signal is also sampled onto the corresponding participant’s surfaces using mri_- vol2surf (FreeSurfer), when surface reconstruction is being performed. Thus, these sampled surfaces can easily be transformed onto different output spaces available by concatenating transforms calculated throughout *fMRIPrep* and internal mappings between spaces calculated with recon-all. The composition of transforms allows for a single-interpolation resampling of volumes using antsApplyTransforms (ANTs). Lanczos interpolation is applied to minimize the smoothing effects of linear or Gaussian kernels^39^. Optionally, ICA-AROMA can be performed and corresponding “non-aggressively” denoised runs are then produced. When ICA-AROMA is enabled, the time-series are first smoothed and then denoised, following the description of the original method^11^.

#### Extraction of nuisance time-series

To avoid restricting *fMRIPrep’s* outputs to particular analysis types, the tool does not perform any temporal denoising by default. Nonetheless, it provides researchers with a diverse set of confound estimates that could be used for explicit nuisance regression or as part of higher-level models. This lends itself to decoupling preprocessing and behavioral modeling as well as evaluating robustness of final results across different denoising schemes. A set of physiological noise regressors are extracted for the purpose of performing component-based noise correction *(CompCor*^10^). Principal components are estimated after high-pass filtering the BOLD time-series (using a discrete cosine filter with 128s cut-off) for the two CompCor variants: temporal (tCompCor) and anatomical (aCompCor). Six tCompCor components are then calculated from the top 5% variable voxels within a mask covering the subcortical regions. This subcortical mask is obtained by heavily eroding the brain mask, which ensures it does not include cortical GM regions. For aCompCor, six components are calculated within the intersection of the aforementioned mask and the union of CSF and WM masks calculated in T1w space, after their projection to the native space of each functional run (using the inverse BOLD-to-T1w transformation). Framewise displacement^40^ and DVARS are calculated for each functional run, both using their implementations in Nipype (following the definitions by Power et al.^7^). Three global signals are extracted within the CSF, the WM, and the whole-brain masks using Nilearn^16^. If ICA-AROMA^11^ is requested, the “aggressive” noise-regressors are collected and placed within the corresponding confounds files. Since the non-aggressive cleaning with ICA-AROMA is performed after extraction of other nuisance signals, the “aggressive” regressors can be used to orthogonalize those other nuisance signals to avoid the risk of re-introducing nuisance signal within regression. In addition, a “non-aggressive” version of preprocessed data is also provided since this variant of ICA-AROMA denoising cannot be performed using only nuisance regressors.

### Visual reports ease quality control and maximize transparency

Users can assess the quality of preprocessing with an individual report generated per participant.Figure 2 shows an example of such reports and describes their structure. Reports contain dynamic and static mosaic views of images at different quality control points along the preprocessing pipeline. Many visual elements of the reports, as well as some of the figures in this manuscript are generated using Nilearn^16^. Only a web browser is required to open the reports on any platform, since they are written in hypertext markup language (HTML). HTML also enables the trivial integration within online neuroimaging services such as OpenNeuro.org, and maximizes shareability between peers. These reports effectively minimize the amount of time required for assessing the quality of the results. They also help understand the internals of processing by visually reporting the full provenance of data throughout the workflow. As an additional transparency enhancement, reports are accompanied by a *citation boilerplate* (see Online Methods, *Box S1)* that follows the guidelines for reporting fMRI studies by Poldrack et al.^41^. Meant for its inclusion within the methodological section of papers using *fMRIPrep*, the boilerplate provides a literate description of the processing that includes software versions of all tools involved in the particular workflow and gives due credit to all authors of all of the individual pieces of software used within *fMRIPrep*.

### Highlights of *fMRIPrep* within the neuroimaging context

*FMRIPrep* is not the first preprocessing pipeline for fMRI data. The most widely used neuroimaging packages generally provide workflows, such as afni_proc.py (AFNI) or `feat` (FSL). Other alternatives include C-PAC^42^ (configurable pipeline for the analysis of connectomes), HCP Pipelines^22^ or the Batch Editor of SPM. In this section, we highlight some additional features beyond robustness and quality that will likely incline scientists to find in *fMRIPrep* the best fit for their fMRI preprocessing needs.

#### Analysis-agnostic: *fMRIPrep* supports a wide range of existing, higher-level analysis and modeling

Alternative workflows are not *agnostic* to currently-available analysis options because they prescribe particular methodologies to analyze the preprocessed data. Important limitations to compatibility with downstream analysis derive from the coordinates space of the outputs and the regular (volume) vs. irregular (surface) sampling of the BOLD signal. For example, HCP Pipelines supports surface-based analyses on subject or template space. Conversely, C-PAC and `feat` are volume-based only. Although afni_proc.py is volume-based by default, pre-reconstructed surfaces can be manually set for sampling the BOLD signal prior to analysis. *FMRIPrep* allows a multiplicity of output spaces including subject-space and atlases for both volume-based and surface-based analyses. While *fMRIPrep* avoids including processing steps that may limit further analysis (e.g. spatial smoothing), other tools are designed to perform preprocessing that supports specific analysis pipelines. For instance, C-PAC performs several processing steps towards the connectivity analysis of resting-state fMRI.

#### Susceptibility distortion correction (SDC) in the absence of field maps

Many legacy and current human fMRI protocols lack the MR field maps necessary to perform standard methods for SDC. *FMRIPrep* adapts the “fieldmap-less” correction method for diffusion echo-planar imaging (EPI) images introduced by Wang et al.^43^. They propose using the same-subject T1w reference as the *undistorted* target in a nonlinear registration scheme. To maximize the similarity between the T2* contrast of the EPI scan and the reference T1w, the intensities of the latter are inverted. To regularize the optimization of the deformation field only displacements along the phase-encoding (PE) direction are allowed, and the magnitude of the displacements is modulated using priors. To our knowledge, no other existing pipeline applies “fieldmap-less” SDC to the BOLD images.

#### FMRIPrep is thoroughly documented, community-driven, and developed with high-standards of software engineering

Preprocessing pipelines are generally well documented, however the extreme flexibility of *fMRIPrep* makes its proper documentation substantially more challenging. As for other large scientific software communities, *fMRIPrep* contributors pledge to keep the documentation thorough and updated along coding iterations. Packages also differ on the involvement of the community: while *fMRIPrep* includes researchers in the decision making process and invites their suggestions and contributions, other packages have a more closed model where the feedback from users is more limited (e.g. a mailing list). In contrast to other pipelines, *fMRIPrep* is community-driven. This paradigm allows the fast adoption of cutting-edge advances on fMRI preprocessing, tend to render existing workflows (including *fMRIPrep)* obsolete. For example, while *fMRIPrep* initially performed STC before HMC, we adapted the tool to the recent recommendations of Power et al.^18^ upon a user’s request*. This model has allowed the user base to grow rapidly and enabled substantial third-party contributions to be included in the software, such as the support for processing datasets without anatomical information. The open-source nature of *fMRIPrep* has permitted frequent code reviews that are effective in enhancing the software’s quality and reliability^44^. Online Methods, sec. Community-centered, peer-reviewed development, and adoption of fMRIPrep describes how the community interacts, discusses the code review process, and underscores how the modular design of *fMRIPrep* successfully facilitates contributions from peers. Finally, *fMRIPrep* undergoes continuous integration testing (see Online Methods, *Figure* S5), a technique that has recently been proposed as a means to ensure reproducibility of analyses in computational sciences^45,46^.

**Figure 2.**
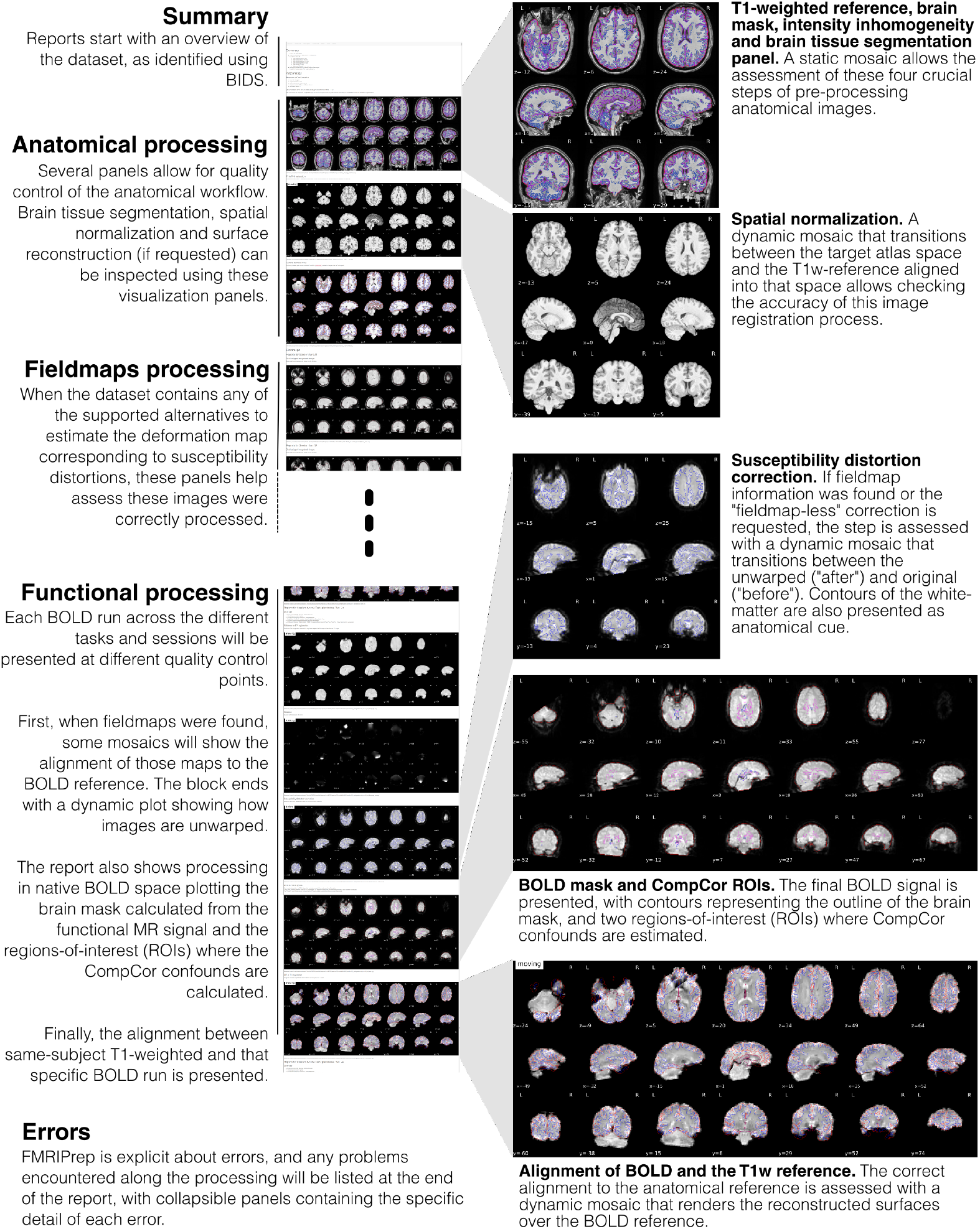
The visual reports ease quality control and help understand the processing flow. The reports enhance the transparency of the tool, because they reflect the multiple steps that the workflow performs, and how these elements are intertwined.

#### Ensuring reproducibility with hard versioning and containers

For enhanced reproducibility, *fMRIPrep* fully supports execution via the Docker (https://docker.com) and Singularity^47^ container platforms. Container images are generated and uploaded to a public repository for each new version of *fMRIPrep*. This helps address the widespread lack of reporting of specific software versions and the large variability of software versions, which threaten the reproducibility of fMRI analyses^19^. These containers are released with a fixed set of software versions for *fMRIPrep* and all its dependencies, maximizing run-to-run reproducibility in an easy way. Except for C-PAC, alternative pipelines do not provide official support for containers. The adoption of the BIDS-Apps^45^ container model makes *fMRIPrep* amenable to a multiplicity of infrastructures and platforms: PC, high-performance computing (HPC), Cloud, etc.

### FMRIPrep yields high-quality results on a diverse set of input data

Figure 3 presents the validation framework that we applied to iteratively maximize the robustness of the tool and validate the quality of the results. The validation framework implements a testing plan elaborated prior the release of the version 1.0 of the software (see Online Methods, sec. Evaluation of *fMRIPrep*). The plan is divided in two validation phases in which different data samples and validation procedures are applied. Table 2 describes the data samples used on each phase and emphasizes how these data are collected from a large number of different, unrelated studies. In Phase I, we ran *fMRIPrep* on a manually selected sample of participants that are potentially challenging to the tool’s robustness, exercising the adaptiveness to the input data. Phase II focused on the visual assessment of the quality of preprocessing results on a large and heterogeneous sample.

**Table 2.** Data from OpenfMRI used in evaluation. S: number of sessions; T: number of tasks; R: number of BOLD runs; Modalities: number of runs for each modality, per subject (FM indicates acquisitions for susceptibility distortion correction); Part. IDs (phase): participant identifiers included in testing phase; N: total of unique participants; TR: repetition time (s); #TR: length of time-series (volumes); Resolution: voxel size of BOLD series (mm).

| DS000XXX | Scanner | S | T | R | Modalities | Part. IDs (Phase I) | Part. IDs (Phase II) | N | TR | #TR | Resolution |
| --- | --- | --- | --- | --- | --- | --- | --- | --- | --- | --- | --- |
| 001<sup>48</sup> | SIEMENS | 1 | 1 | 21 | 1 T1w, 3 BOLD | 02, 03, 09, 15 | 01, 02, 07, 08 | 7 | 2.0 | 6300 | 3.12x3.12x4.00 |
| 002<sup>49</sup> | SIEMENS | 1 | 3 | 48 | 1 T1w, 6 BOLD | 01, 11, 14, 15 | 02, 03, 04, 10 | 8 | 2.0 | 9510 | 3.12x3.12x5.00 |
| 003<sup>50</sup> | SIEMENS | 1 | 1 | 6 | 1 T1w, 1 BOLD | 03, 07, 09, 11 | 02, 09, 10, 11 | 6 | 2.0 | 956 | 3.12x3.12x4.00 |
| 005<sup>51</sup> | SIEMENS | 1 | 1 | 21 | 1 T1w, 3 BOLD | 01, 03, 06, 14 | 01, 04, 05, 15 | 7 | 2.0 | 5040 | 3.12x3.12x4.00 |
| 007<sup>52</sup> | SIEMENS | 1 | 3 | 46 | 1 T1w, 5 BOLD | 09, 11, 18, 20 | 03, 04, 08, 12 | 8 | 2.0 | 8205 | 3.12x3.12x4.00 |
| 008<sup>53</sup> | SIEMENS | 1 | 2 | 38 | 1 T1w, 5 BOLD | 04, 09, 12, 14 | 10, 12, 13, 15 | 7 | 2.0 | 6808 | 3.12x3.12x4.39 |
| 009 | SIEMENS | 1 | 4 | 48 | 1 T1w, 6 BOLD | 01, 03, 09, 10 | 17, 18, 21, 23 | 8 | 2.0 | 10528 | 3.00x3.00x4.00 |
| 011<sup>54</sup> | SIEMENS | 1 | 4 | 41 | 1 T1w, 5 BOLD | 01, 03, 06, 08 | 03, 09, 11, 14 | 7 | 2.0 | 8041 | 3.12x3.12x5.00 |
| 017 | SIEMENS | 2 | 2 | 48 | 4 T1w, 9 BOLD | 2, 4, 7, 8 | 2, 5, 7, 8 | 5 | 2.0 | 8736 | 3.12x3.12x4.00 |
| 030<sup>55,56</sup> | SIEMENS | 1 | 8 | 30 | 1 T1w, 7 BOLD |  | 10[440,638,668,855] | 4 | 2.2 | 6254 | 3.00x3.00x4.00 |
| 031<sup>57</sup> | SIEMENS | 107 | 9 | 191 | 29 T1w, 18 T2w, 46 FM, 191 BOLD |  | 01 | 1 | 1.2 | 79017 | 2.55x2.55x2.54 |
| 051<sup>58</sup> | SIEMENS | 1 | 1 | 54 | 2 T1w, 7 BOLD | 03, 04, 05, 13 | 02, 04, 06, 09 | 7 | 2.0 | 10800 | 3.12x3.12x6.00 |
| 052<sup>59</sup> | SIEMENS | 1 | 2 | 28 | 2 T1w, 4 BOLD | 06, 08, 12, 14 | 05, 10, 12, 13 | 7 | 2.0 | 6300 | 3.12x3.12x6.00 |
| 053 | SIEMENS | 1 | 3 | 32 | 1 T1w, 8 BOLD |  | 002, 003, 005, 006 | 4 | 1.2 | 10712 | 2.40x2.40x2.40 |
| 101 | SIEMENS | 1 | 1 | 16 | 1 T1w, 2 BOLD | 06, 08, 16, 19 | 05, 11, 17, 20 | 8 | 2.0 | 2416 | 3.00x3.00x4.00 |
| 102<sup>60-62</sup> | SIEMENS | 1 | 1 | 16 | 1 T1w, 2 BOLD | 05, 19, 22, 23 | 08, 10, 16, 20 | 8 | 2.0 | 2336 | 3.00x3.00x4.00 |
| 105<sup>63,64</sup> | GE | 1 | 1 | 71 | 1 T1w, 11 BOLD | 1, 2, 3, 6 | 1, 4, 5, 6 | 6 | 2.5 | 8591 | 3.50x3.75x3.75 |
| 107<sup>65</sup> | SIEMENS | 1 | 1 | 14 | 1 T1w, 2 BOLD | 02, 05, 20, 29 | 05, 36, 39, 47 | 7 | 3.0 | 2315 | 3.00x3.00x3.00 |
| 108<sup>66</sup> | GE | 1 | 1 | 41 | 1 T1w, 5 BOLD | 01, 03, 07, 17 | 03, 10, 24, 26 | 7 | 2.0 | 7860 | 3.44x3.44x4.50 |
| 109<sup>67</sup> | SIEMENS | 1 | 1 | 12 | 1 T1w, 2 BOLD | 02, 10, 39, 47 | 02, 11, 15, 39 | 6 | 2.0 | 2148 | 3.00x3.00x3.54 |
| 110<sup>68</sup> | GE | 1 | 1 | 80 | 1 T1w, 10 BOLD | 07, 09, 17, 18 | 01, 02, 03, 06 | 8 | 2.0 | 14880 | 3.44x3.44x4.01 |
| 114<sup>69</sup> | GE | 2 | 5 | 70 | 2 T1w, 10 BOLD | 01, 05, 07, 08 | 02, 03, 04, 07 | 7 | 5.0 | 10626 | 4.00x4.00x4.00 |
| 115<sup>70,71</sup> | SIEMENS | 1 | 3 | 24 | 1 T1w, 3 BOLD | 31, 68, 77, 78 | 04, 33, 67, 79 | 8 | 2.5 | 3288 | 4.00x4.00x4.00 |
| 116<sup>72-75</sup> | PHILIPS | 1 | 2 | 36 | 1 T1w, 6 BOLD | 02, 08, 10, 15 | 08, 12, 15, 17 | 6 | 2.0 | 6120 | 3.00x3.00x4.00 |
| 119<sup>76</sup> | SIEMENS | 1 | 1 | 31 | 1 T1w, 3 BOLD | 10, 51, 59, 74 | 11, 26, 56, 58 | 8 | 1.5 | 7564 | 3.12x3.12x4.00 |
| 120<sup>77</sup> | SIEMENS | 1 | 1 | 11 | 1 T1w, 2 BOLD |  | 04, 05, 08, 24 | 4 | 1.5 | 2376 | 3.12x3.12x4.00 |
| 121<sup>78</sup> | SIEMENS | 1 | 1 | 28 | 1 T1w, 4 BOLD | 01, 04, 05, 20 | 01, 18, 22, 26 | 7 | 1.5 | 5656 | 3.12x3.12x4.00 |
| 133<sup>79</sup> | PHILIPS | 2 | 1 | 24 | 2 T1w, 6 BOLD |  | 06, 21, 22, 23 | 4 | 1.7 | 3480 | 4.00x4.00x4.00 |
| 140<sup>80</sup> | PHILIPS | 1 | 1 | 36 | 1 T1w, 9 BOLD |  | 05, 27, 32, 33 | 4 | 2.0 | 7380 | 2.80x2.80x3.00 |
| 148 | GE | 1 | 1 | 12 | 1 T1w, 1 T2w, 3 BOLD |  | 09, 26, 28, 33 | 4 | 1.8 | 3162 | 3.00x3.00x3.00 |
| 157<sup>81</sup> | PHILIPS | 1 | 1 | 4 | 1 T1w, 1 BOLD |  | 04, 21, 23, 28 | 4 | 1.6 | 1485 | 4.00x4.00x3.99 |
| 158<sup>82</sup> | SIEMENS | 1 | 1 | 4 | 1 T1w, 1 BOLD |  | 064, 081, 122, 149 | 4 | 2.0 | 1240 | 3.00x3.00x3.30 |
| 164<sup>83</sup> | SIEMENS | 1 | 1 | 4 | 1 T1w, 1 BOLD |  | 006, 012, 019, 027 | 4 | 1.5 | 1480 | 3.50x3.50x3.50 |
| 168<sup>84</sup> | SIEMENS | 1 | 1 | 4 | 1 T1w, 1 BOLD |  | 08, 27, 30, 49 | 4 | 2.5 | 2112 | 3.00x3.00x3.00 |
| 170<sup>85-87</sup> | GE | 1 | 4 | 48 | 1 T1w, 12 BOLD |  | 1700, 1708, 1710, 1713 | 4 | 3.0 | 2160 | 3.44x3.44x3.40 |
| 171<sup>88</sup> | SIEMENS | 1 | 2 | 20 | 1 T1w, 5 BOLD |  | control0[4,8,14], mdd03 | 4 | 3.0 | 2066 | 2.90x2.90x3.00 |
| 177<sup>89</sup> | SIEMENS | 1 | 1 | 4 | 1 T1w, 1 BOLD |  | 04, 07, 10, 11 | 4 | 3.0 | 920 | 3.00x3.00x3.00 |
| 200<sup>90</sup> | SIEMENS | 1 | 1 | 4 | 1 T1w, 1 BOLD |  | 2004, 2011, 2012, 2014 | 4 | 2.5 | 480 | 3.28x3.28x4.29 |
| 205<sup>91</sup> | SIEMENS | 1 | 2 | 12 | 1 T1w, 3 BOLD |  | 01, 05, 06, 07 | 4 | 2.2 | 4103 | 3.00x3.00x3.00 |
| 208<sup>92</sup> | SIEMENS | 1 | 1 | 4 | 1 T1w, 1 BOLD |  | 27, 45, 56, 69 | 4 | 2.5 | 1200 | 3.44x3.44x3.00 |
| 212<sup>93,94</sup> | SIEMENS | 1 | 2 | 40 | 1 T1w, 10 BOLD |  | 07, 13, 20, 29 | 4 | 3.0 | 5808 | 3.12x3.12x4.00 |
| 213<sup>95</sup> | SIEMENS | 1 | 1 | 4 | 1 T1w, 1 BOLD |  | 06, 10, 12, 13 | 4 | 2.0 | 1120 | 3.00x3.00x3.99 |
| 214<sup>96</sup> | SIEMENS | 1 | 1 | 4 | 1 T1w, 1 BOLD |  | EES0[06,31,33,34] | 4 | 1.6 | 1364 | 3.44x3.44x5.00 |
| 216<sup>97</sup> | GE | 1 | 1 | 16 | 1 T1w, 4 BOLD (ME) |  | 01, 02, 03, 04 | 4 | 3.5 | 2688 | 3.00x3.00x3.00 |
| 218<sup>98</sup> | PHILIPS | 1 | 1 | 12 | 1 T1w, 3 BOLD |  | 02, 07, 12, 17 | 4 | 1.5 | 6709 | 2.88x3.00x2.88 |
| 219<sup>98</sup> | PHILIPS | 1 | 1 | 14 | 1 T1w, 3 BOLD |  | 04, 09, 10, 12 | 4 | 1.5 | 7807 | 2.88x3.00x2.88 |
| 220<sup>99</sup> | PHILIPS, SIEMENS | 3 | 1 | 12 | 3 T1w, 3 BOLD |  | tbi[03,05,06,10] | 4 | 2.0 | 1728 | 3.00x3.00x4.00 |
| 221 | SIEMENS | 2 | 1 | 15 | 1 MP2RAGE, 9 FM, 3 BOLD |  | 010[016,064,125,251] | 4 | 2.5 | 9855 | 2.30x2.30x2.30 |
| 224<sup>100</sup> | SIEMENS | 12 | 6 | 399 | 4 T1w, 4 T2w, 10 FM, 79 BOLD | MSC[05,06,08,09] | MSC[05,08,09,10] | 5 | 2.2 | 88528 | 4.00x4.00x4.00 |
| 228 | SIEMENS | 1 | 1 | 4 | 1 T1w, 1 BOLD |  | pixar[001,017,103,132] | 4 | 2.0 | 672 | 3.06x3.06x3.29 |
| 229<sup>101</sup> | SIEMENS | 1 | 1 | 12 | 1 T1w, 3 BOLD |  | 02, 05, 07, 10 | 4 | 2.0 | 4680 | 3.44x3.44x3.00 |
| 231<sup>102</sup> | SIEMENS | 1 | 1 | 12 | 1 T1w, 3 BOLD |  | 01, 02, 03, 09 | 4 | 2.0 | 4548 | 2.02x2.02x2.00 |
| 233<sup>103</sup> | PHILIPS | 1 | 2 | 80 | 2 T1w, 10 BOLD | rid0000[12,24,36,41] | rid0000[01,17,31,32] | 8 | 2.0 | 15680 | 3.00x3.00x3.00 |
| 237<sup>104</sup> | SIEMENS | 1 | 1 | 41 | 1 T1w, 5 BOLD | 03, 08, 11, 12 | 01, 03, 04, 06 | 7 | 1.0 | 19844 | 3.00x3.00x3.00 |
| 243<sup>40</sup> | SIEMENS | 1 | 1 | 13 | 1 T1w, 1 BOLD | 012, 032, 042, 071 | 023, 066, 089, 094 | 8 | 2.5 | 2884 | 4.00x4.00x4.00 |
| Total |  |  |  | 2176 |  | 120 | 202 | 304 |  | 551769 |  |

#### Validation Phase I – Fault-discovery testing

We tested *fMRIPrep* on a set of 30 datasets from OpenfMRI (see Table 2). Included participants were manually selected for their low quality as visually assessed by two experts using MRIQC^105^ (the assessment protocol is further described in in Online Methods, sec. Evaluation of *fMRIPrep*). Data showing substandard quality are known to likely degrade the outcomes of image processing^105^, and therefore they are helpful to test software reliability. Phase I concluded with the release of *fMRIPrep* version 1.0 on December 6, 2017.

#### Validation Phase II – Quality assurance and reliability testing

We extended the evaluation data up to 54 datasets from OpenfMRI (see Table 2). Participants were selected randomly as described in Online Methods, sec. Evaluation of *fMRIPrep*. Validation Phase II integrated a protocol for the screening of results into the software testing (Figure 3). As shown in Figure 4, this effectively contributed to substantive improvements on the quality of results. Three raters (authors CJM, KJG and OE) evaluated the 213 visual reports at six quality control points throughout the pipeline, and also assigned an overall score to each participant. Their ratings are made available with the corresponding reports for scrutiny. The scoring scale has three levels: 1 (“poor”), 2 (“acceptable”) and 3 (“excellent”). A special rating of 0 (“unusable”) is assigned to critical failures that hamper any further processing beyond the quality control checkpoint. Further details on how these quality categories were assigned are given in Online Methods, sec. Iterative quality and robustness assurance. After Phase II, 50 datasets out of the total 54 were rated above the “acceptable” average quality level. The remaining 4 datasets were all above the “poor” level and in or nearby the “acceptable” rating. Figure 4 illustrates the quality of results, while Online Methods, *Figure S6* shows the individual evolution of every dataset at each of the seven quality control points. Phase II concluded with the release of *fMRIPrep* version 1.0.8 on February 22, 2018.

**Figure 3.**
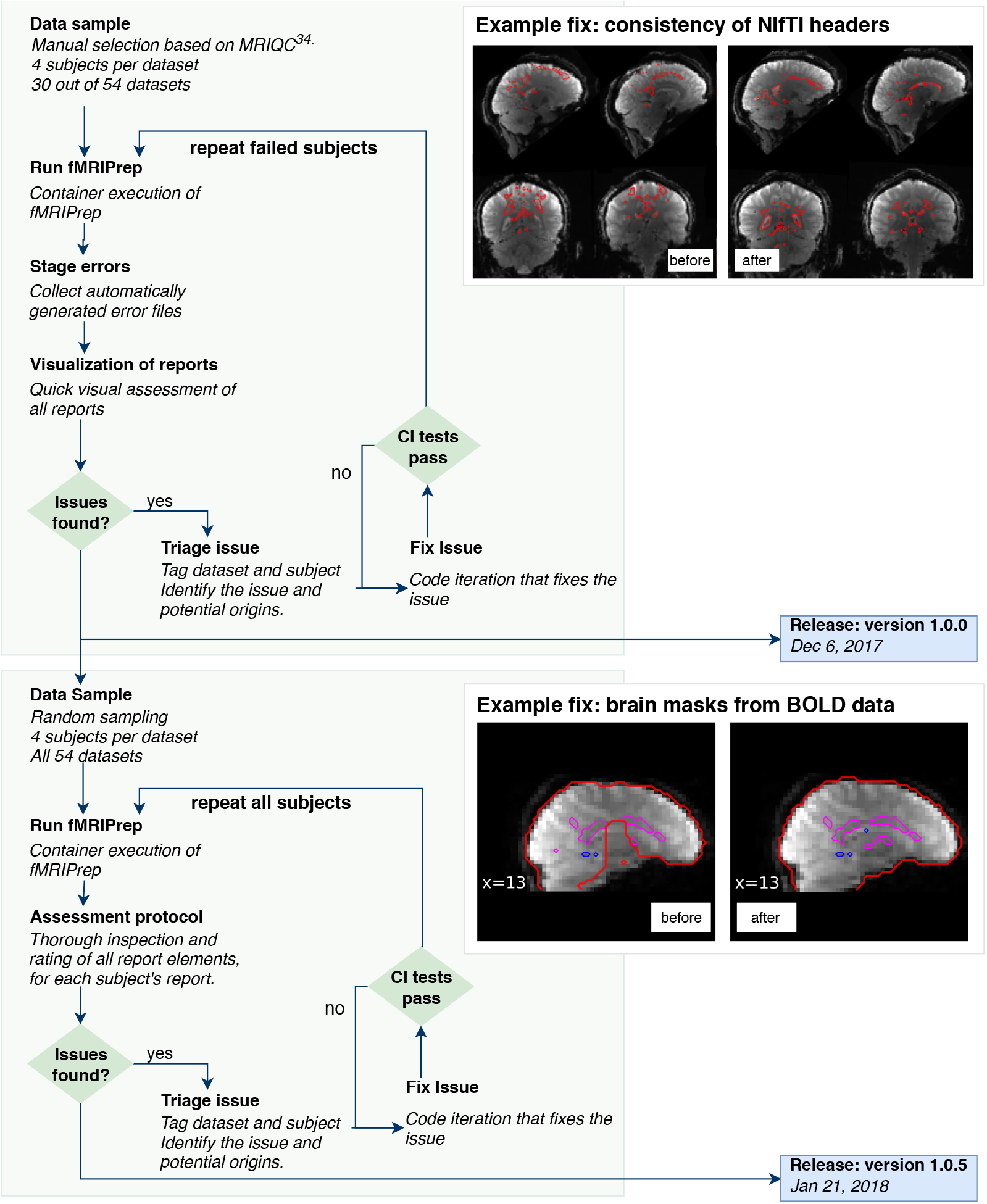
Combining visual assessment within the software testing flow. We complement well-established techniques for software integration testing with manual assessment of the outputs. The evaluation framework is designed with two subsequent testing phases. Phase I focuses on fault-discovery and visual reports are used to better understand the issues found. The top box (Example fix 1) shows an example of defect identified and solved during this testing cycle. After addressing a total of 21 issues affecting 7 datasets, and the release of *fMRIPrep* version 1.0.0, the next testing stage is initiated. Phase II focuses on increasing the overall quality of results as evaluated visually by experts. Following an inspection protocol, reports from 213 participants belonging to 58 different studies were individually assessed. We found 12 additional issues affecting 11 datasets that have been addressed with the release of *fMRIPrep* version 1.0.3 on January 3, 2018. The bottom box (Example fix 2) illustrates one of these issues, which produced errors in the brain extraction process from BOLD data. In Online Methods, sec. Iterative quality and robustness assurance, both examples and the corresponding solutions are described in detail.

**Figure 4.**
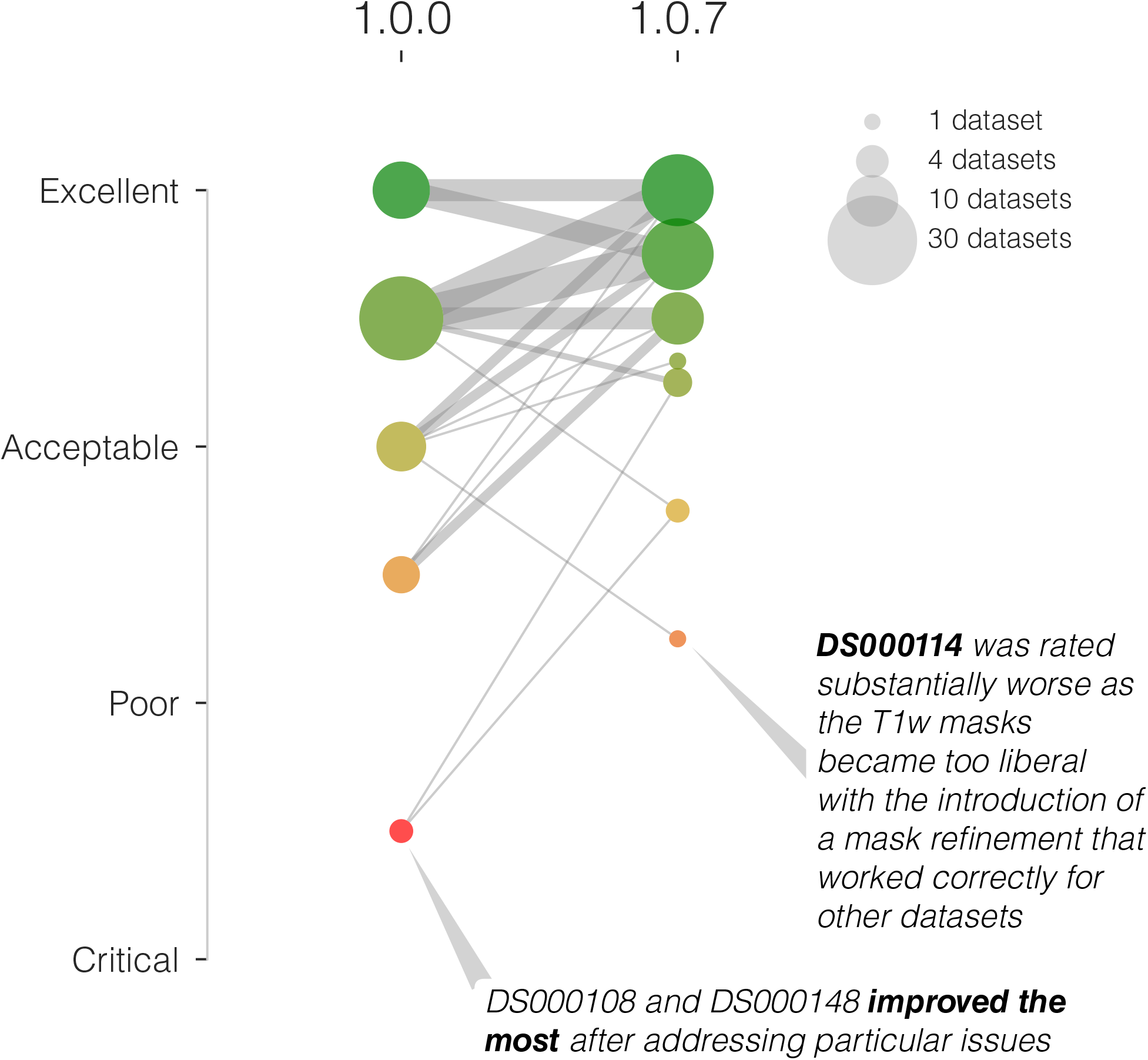
Integrating visual assessment into the software testing framework effectively increases the quality of results. In an early assessment of quality using *fMRIPrep* version 1.0.0, the overall rating of two datasets was below the “poor” category and four below the “acceptable” level (left column of colored circles). After addressing some outstanding issues detected by the early assessment, the overall quality of processing is substantially improved (right column of circles), and no datasets are below the “poor” quality level. Only two datasets are rated below the “acceptable” level in the second assessment (using *fMRIPrep* version 1.0.7).

### FMRIPrep improves spatial precision through reduced smoothing

We investigate whether the focus on robustness against data irregularity comes at a cost in quality of the preprocessing outcomes by comparing it to the commonly used FSL `feat` workflow. Using all the scans of the “stopsignal” task in DS000030 (N=257 participants) from OpenfMRI, we ran *fMRIPrep* and a standard `feat` workflow. We chose `feat` because DS000030 had successfully been preprocessed and analyzed with FSL tools previously^55^. Smoothing is intentionally excluded from both preprocessing routes with the aim to apply it early within a common (identical) analysis workflow. We calculated standard deviation maps in MNI space^106^ for the temporal average map of the “stopsignal” task derived from preprocessing with both alternatives. Visual inspection of these variability maps (Figure 5) reveals a higher anatomical accuracy of *fMRIPrep* over `feat`, likely reflecting the combined effects of a more precise spatial normalization scheme and the application of “fieldmap-less” SDC. *FMRIPrep* outcomes are particularly better aligned with the underlying anatomy in regions typically warped by susceptibility distortions such as the orbitofrontal lobe, as demonstrated by close-ups in Online Methods, *Figure S7*.

We also compared preprocessing done with *fMRIPrep* and FSL’s `feat` in two common fMRI analyses. First, we performed within subject statistical analysis using `feat` –the same tool provides preprocessing and first-level analysis– on both sets of preprocessed data. Second, we perform a group statistical analysis using ordinary least squares (OLS) mixed modeling (`flame`^107^, FSL). In both experiments, we applied identical analysis workflows and settings to both preprocessing alternatives. Using AFNI’s 3dFWHMx, we estimated the smoothness of data right after preprocessing (unsmoothed), and after an initial smoothing step of 5.0mm (full-width half-minimum, FWHM) of the common analysis workflow. As visually suggested by Figure 5, we indeed found that `feat` produces smoother data (Figure 6A). Although preprocessed data were resampled to an isotropic voxel size of 2.0×2.0×2.0 [mm], the smoothness estimation (before the prescribed smoothing step) for *fMRIPrep* was below 4.0mm, very close to the original resolution of 3.0×3.0×4.0 [mm] of these data. The first-level analysis showed that the thresholded activation count maps for the go vs. successful stop contrast in the “stopsignal” task were very similar *(Figure* 6B). It can be seen that the results from both pipelines identified activation in the same regions. However, since data preprocessed with `feat` are smoother, the results from *fMRIPrep* are more local and better aligned with the cortical sheet.

**Figure 5.**
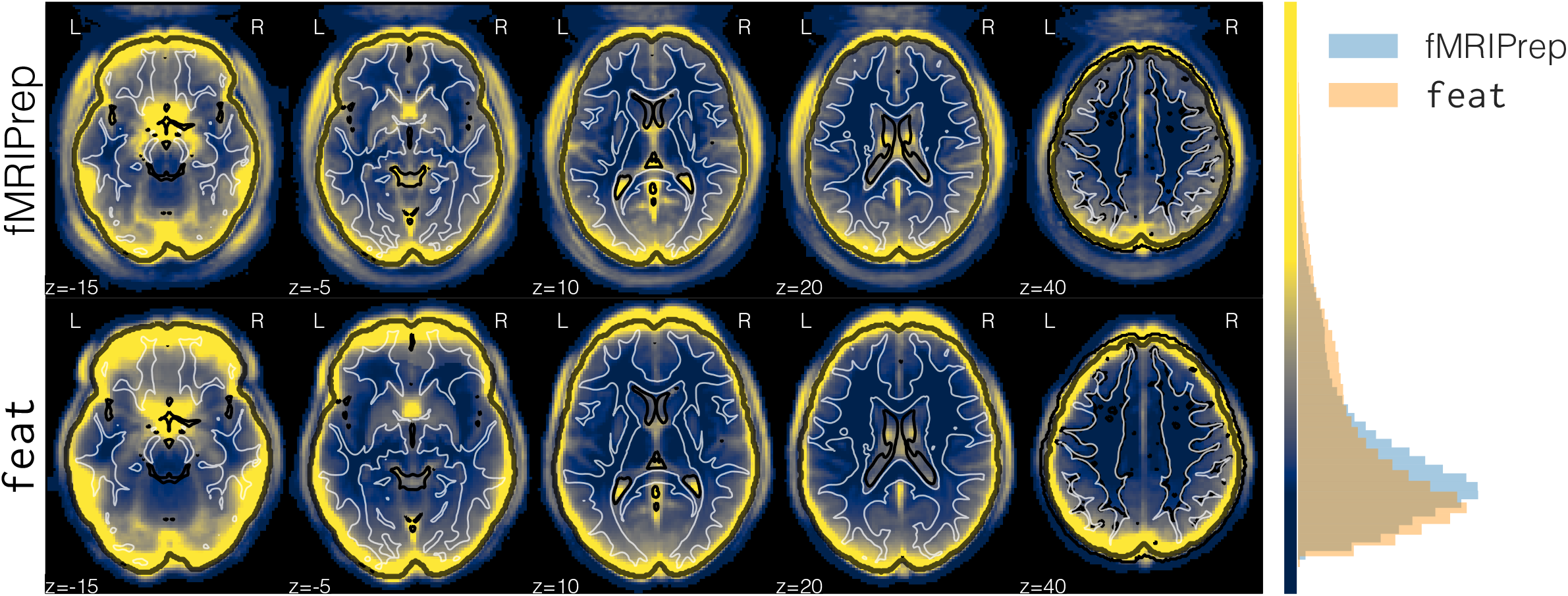
Maps of between-subjects variability of the averaged BOLD time-series resampled into MNI space. We preprocessed DS000030 (N=257) with *fMRIPrep* and FSL feat. This figure shows greater between-subject variability of the averaged BOLD series obtained with feat, in MNI space. The top box of the panel shows these maps at different axial planes of the image grid, with reference contours from the MNI atlas. The map summarizing feat-derived results displays greater variability around the brain outline (represented with a black contour). This effect is generally associated with a lower performance of spatial normalization^106^. The histogram at the right side plots the normalized frequency of variability (arbitrary units) for both maps, within the brain mask. The color bar maps the bins of the histogram to the corresponding colors that are used in the left panel. The heavier tail of `feat` in the distribution of variability is compensated in the case of *fMRIPrep* by the peak in voxel-counts located at a higher variability than that of feat. Such peak in *fMRIPrep*’s distribution corresponds to variability in cortical gray matter regions. See Online Methods, *Figure S7* for close-ups into regions affected by susceptibility-derived distortions.

To investigate the implications of either pipeline on the group analysis use-case, we run the same OLS modeling on two disjoint subsets of randomly selected subjects. We calculate several metrics of spatial agreement on the resulting maps of (uncorrected) *p*-statistical values, and also after binarizing these maps with a threshold chosen to control for the false discovery rate at 5%. The overlap of statistical maps, as well as Pearson’s correlation, were tightly related to the smoothing of the input data. In Online Methods, sec. Comparison to FSL `feat` we report the group-level analysis in full. We ran two variants of the analysis: with a prescribed smoothing of 5.0mm FWHM, and without the smoothing step. These results showed that, at the group-level analysis, *fMRIPrep* and `feat` perform equivalently.

## DISCUSSION

*FMRIPrep* is an fMRI preprocessing workflow developed to excel at four aspects of scientific software: *robustness* to data idiosyncrasies, high *quality* and consistency of results, maximal *transparency*, and *ease-of-use*. We describe how using the Brain Imaging Data Structure (BIDS^24^) along with a flexible design allows the workflow to self-adapt to the idiosyncrasy of inputs (sec. A modular design allows for a flexible, adaptive workflow). The workflow (briefly summarized in Figure 1) integrates state-of-art tools from widely used neuroimaging software packages at each preprocessing step (see Table 1). Some other relevant facets of *fMRIPrep* and how they relate to existing alternative pipelines are presented in sec. Highlights of *fMRIPrep* within the neuroimaging context. We stress that *fMRIPrep* is developed with the best software engineering principles, which are fundamental to ensure software reliability. The pipeline is easy to use for researchers and clinicians without extensive computer engineering experience, and produces comprehensive visual reports *(Figure* 2).

In sec. *FMRIPrep* yields high-quality results on a diverse set of input data, we demonstrate the robustness of *fMRIPrep* on a representative collection of data from datasets associated with different studies (Table 2). We then interrogate the quality of those results with the individual inspection of the corresponding visual reports by experts (sec. Visual reports ease quality control and maximize transparency and the corresponding summary in Figure 4). A comparison to FSL’s `feat` (sec. *FMRIPrep* improves spatial precision through reduced smoothing) demonstrates that *fMRIPrep* achieves higher spatial accuracy and introduces less uncontrolled smoothness (Figures 5, 6). Group *p*-statistical maps only differed on their smoothness (sharper for the case of *fMRIPrep*). The fact that first-level and second-level analyses resulted in small differences between *fMRIPrep* and our *ad hoc* implementation of a feat-based workflow indicates that the individual preprocessing steps perform similarly when they are fine-tuned to the input data. That justifies the need for *fMRIPrep*, which autonomously adapts the workflow to the data without error-prone manual intervention. To a limited extent, that also mitigates some concerns and theoretical risks arisen from the analytical degrees-of-freedom^19^ available to researchers. *FMRIPrep* stands out amongst pipelines because it automates the adaptation to the input dataset without compromising the quality of results.

**Figure 6.**
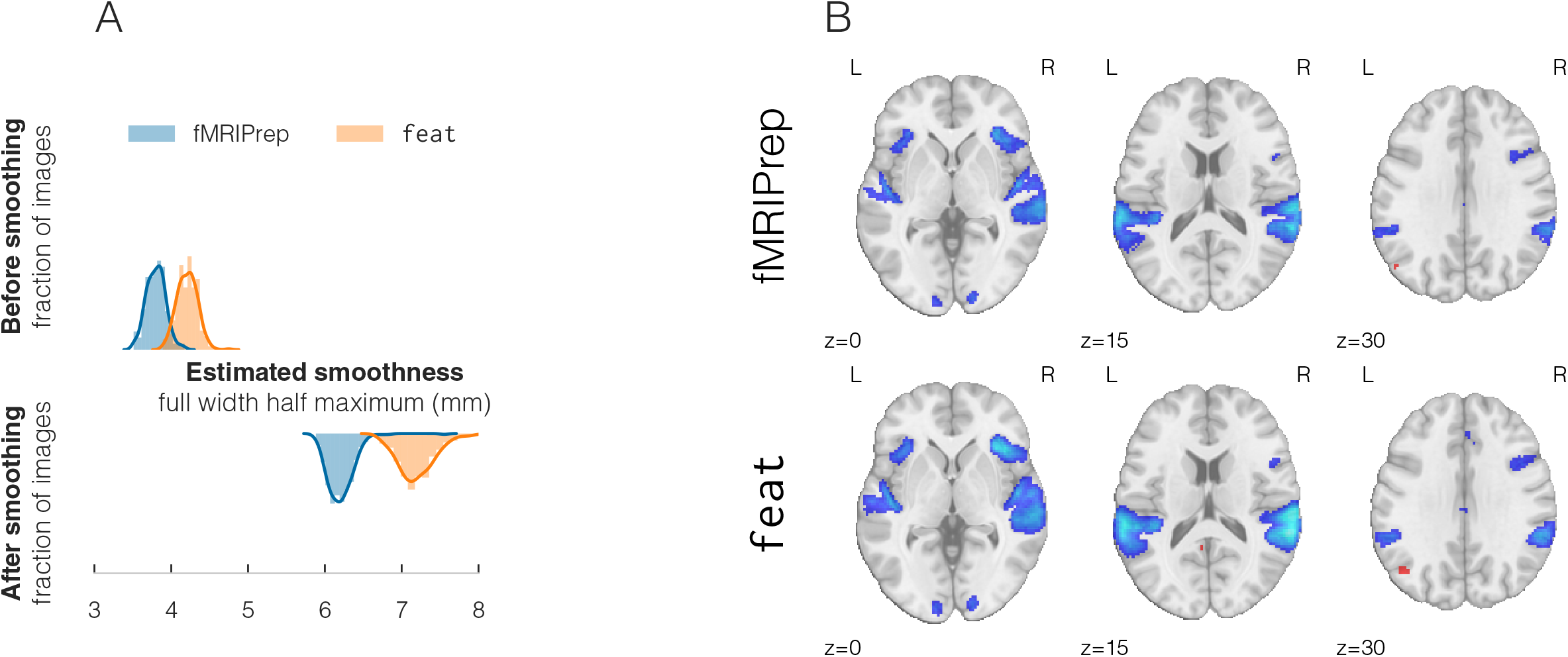
**A** | Estimating the spatial smoothness of data before and after the initial smoothing step of the analysis workflow confirmed that results of preprocessing with `feat` are intrinsically smoother. Therefore, *fMRIPrep* affords the researcher finer control over the smoothness of their analysis. **B** | Thresholded activation count maps for the go vs. successful stop contrast in the “stopsignal” task after preprocessing using either *fMRIPrep* or FSL’s feat, with identical single subject statistical modeling. Both tools obtained similar activation maps, with *fMRIPrep* results being slightly better aligned with the underlying anatomy.

One limitation of this work is the use of visual (the reports) and semi-visual (e.g. Figure 5 and Figure 6) assessments for the quality of preprocessing outcomes. Although some frameworks have been proposed for the quantitative evaluation of preprocessing on task-based (such as NPAIRS^108^) and resting-state^109^ fMRI, they impose a set of assumptions on the test data and the workflow being assessed that severely limit their suitability. The modular design of *fMRIPrep* defines an interface to each processing step, which will permit the programmatic evaluation of the many possible combinations of software tools and processing steps. That will also enable the use of quantitative testing frameworks to pursue the minimization of Type I errors without the cost of increasing Type II errors.

The range of possible applications for *fMRIPrep* also presents some boundaries. For instance, very narrow field-of-view (FoV) images oftentimes do not contain enough information for standard image registration methods to work correctly. Reduced FoV datasets from OpenfMRI were excluded from the evaluation since they are not yet fully supported by *fMRIPrep*. Extending *fMRIPrep’*s support for these particular images is already a future line of the development road-map. *FMRIPrep* may also under-perform for particular populations (e.g. infants) or when brains show nonstandard structures, such as tumors, resected regions or lesions. Nonetheless, *fMRIPrep’s* modular architecture makes it straightforward to extend the tool to support specific populations or new species by providing appropriate atlases of those brains. This future line of work would be particularly interesting in order to adapt the workflow to data collected from rodents and nonhuman primates. By contrast, *fMRIPrep* performed robustly on data from a simultaneous MRI/electrocorticography (ECoG) study, which is extremely challenging to analyze due to the massive BOLD signal drop-out near the implanted cortical electrodes (see Online Methods, *Figure S11*). Likewise, *fMRIPrep* successfully preprocessed multiple multi-band datasets.

Approximately 80% of the analysis pipelines investigated by Carp^19^ were implemented using either AFNI^12^, FSL^15^, or SPM^17^. *Ad hoc* pipelines adapt the basic workflows provided by these tools to the particular dataset at hand. Although workflow frameworks like Nipype^110^ ease the integration of tools from different packages, these pipelines are typically restricted to just one of these alternatives (AFNI, FSL or SPM). Otherwise, scientists can adopt the acquisition protocols and associated preprocessing software of large consortia like the Human Connectome Project (HCP) or the UK Biobank. The *off-the-shelf* applicability of these workflows is contravened by important limitations on the experimental design. Therefore, researchers typically opt to recode their custom preprocessing workflows with nearly every new study^19^. That practice entails a “pipeline debt”, which requires the investment on proper software engineering to ensure an acceptable correctness and stability of the results (e.g. continuous integration testing) and reproducibility (e.g. versioning, packaging, containerization, etc.). A trivial example of this risk would be the leakage of *magic numbers* that are hard-coded in the source (i.e. a crucial imaging parameter that inadvertently changed from one study to the next one). Until *fMRIPrep*, an analysis-agnostic approach that builds upon existing software instruments and optimizes preprocessing for robustness to data idiosyncrasies, quality of outcomes, ease-of-use, and transparency, was lacking.

The rapid increase in volume and diversity of data, as well as the evolution of available techniques for processing and analysis, presents an opportunity for significantly advancing research in neuroscience. The drawback resides in the need for progressively complex analysis workflows that rely on decreasingly interpretable models of the data. Such context encourages “black-box” solutions that efficiently perform a valuable service but do not provide insights into how the tool has transformed the data into the expected outputs. Black-boxes obscure important steps in the inductive process mediating between experimental measurements and reported findings. This way of moving forward risks producing a future generation of cognitive neuroscientists who have become experts in using sophisticated computational methods, but have little to no working knowledge of how data were transformed through processing. Transparency is often identified as a treatment for these problems. *FMRIPrep* ascribes to “glass-box” principles, which are defined in opposition to the many different facets or levels at which black-box solutions are opaque.

The visual reports that *fMRIPrep* generates are a crucial aspect of the glass-box. Their quality control checkpoints represent the logical flow of preprocessing, allowing scientists to critically inspect and better understand the underlying mechanisms of the workflow. A second transparency element is the citation boilerplate that formalizes all details of the workflow and provides the versions of all involved tools along with references to corresponding scientific literature. A third asset for transparency is the thorough documentation which delivers additional details on each of the building blocks that are represented in the visual reports and described in the boilerplate. Further, *fMRIPrep* is open-source since its inception: users have access to all the incremental additions to the tool through the history of the version-control system. The use of GitHub (https://github.com/poldracklab/fmriprep) grants access to the discussions held during development, allowing the retrieval of how and why the main design decisions were made. GitHub also provides an excellent platform to foster the community and provides useful tools such as source browsing, code review, bug tracking and reporting, submission of new features and bug fixes through pull-requests, etc. The modular design of *fMRIPrep* enhances its flexibility and helps transparency, as the main features of the software are more easily accessible to potential collaborators. In combination to some coding style and contribution guidelines, this modularity has enabled multiple contributions by peers and the creation of a rapidly growing community that would be difficult to nurture behind closed doors. A number of existing tools have implemented elements of “glass-box” philosophy (for example visual reports in FEAT, documentation in C-PAC, open source community of Nilearn), but the complete package (visual reports, educational documentation, reporting templates, collaborative open source community) is still rare among scientific software. *FMRIPrep*’s transparent and accessible development and reporting aims to better equip fMRI practitioners to perform reliable, reproducible, statistical analyses with a high-standard, consistent, and adaptive preprocessing instrument.

## CONCLUSION

Despite efforts to achieve high-quality preprocessing of idiosyncratic fMRI datasets, doing so reliably has remained an open problem. *FMRIPrep* is an analysis-agnostic, preprocessing workflow that yields consistent results across a wide range of input datasets. *FMRIPrep* is built on top of the best neuroimaging tools selected from various software packages. These tools are integrated into workflows that can be dynamically combined to compose a full preprocessing workflow adapted to the input data. The optimal workflow for the input dataset is constructed at runtime, blending a set of heuristics with the Brain Imaging Data Structure (BIDS) to read the inputs. *FMRIPrep* excels in four design goals: robustness, high-quality of results, transparency and ease-of-use. To validate and demonstrate these features, we integrate the individual screening of preprocessing results with continuous integration techniques of software testing. The process is aided by comprehensive, portable reports that inform the scientist about the workflow, ease the quality control of results and maximize the shareability of research outcomes. We highlight the aspects that justify the development of *fMRIPrep* with respect to currently available preprocessing workflows. We quantitatively demonstrate that *fMRIPrep* does not introduce uncontrolled smoothing as compared to one alternative software. *FMRIPrep* aims to better equip fMRI practitioners to perform reliable, reproducible statistical analyses with a high-standard, transparent, and verifiable instrument.

## Footnotes

* https://neurostars.org/t/obtaining-movement-estimates-before-slice-time-correction/1007

## ACKNOWLEDGMENTS

This work was supported by the Laura and John Arnold Foundation, NIH R01 EB020740, NIH 1R24MH114705-01, and NINDS grant 1U01NS103780-01. JD has received funding from the European Union’s Horizon 2020 research and innovation program under the Marie Sklodowska-Curie grant agreement No 706561. The authors thank S. Nastase and T. van Mourik for their thoughtful open reviews of a pre-print version of this paper.

---

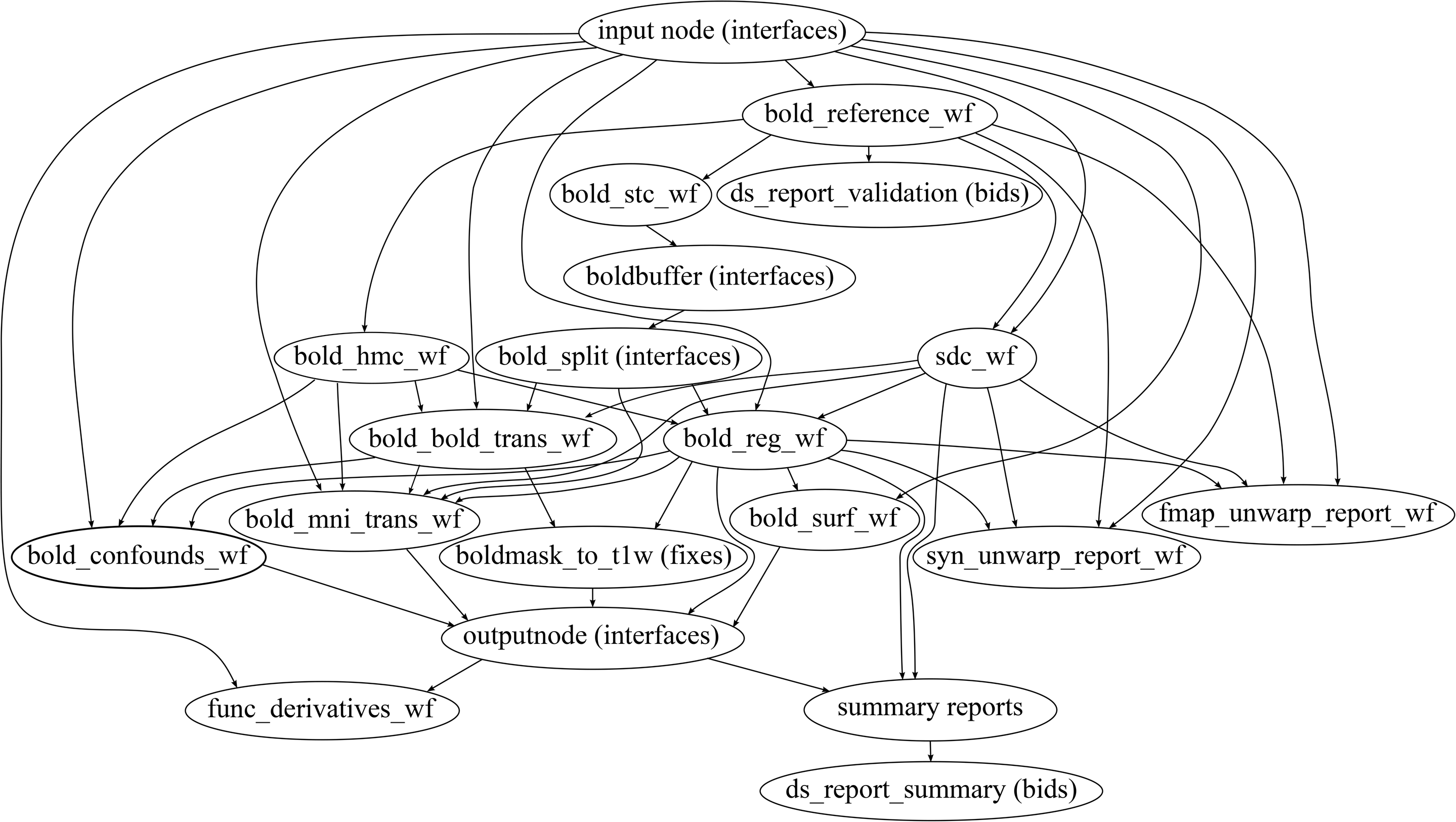

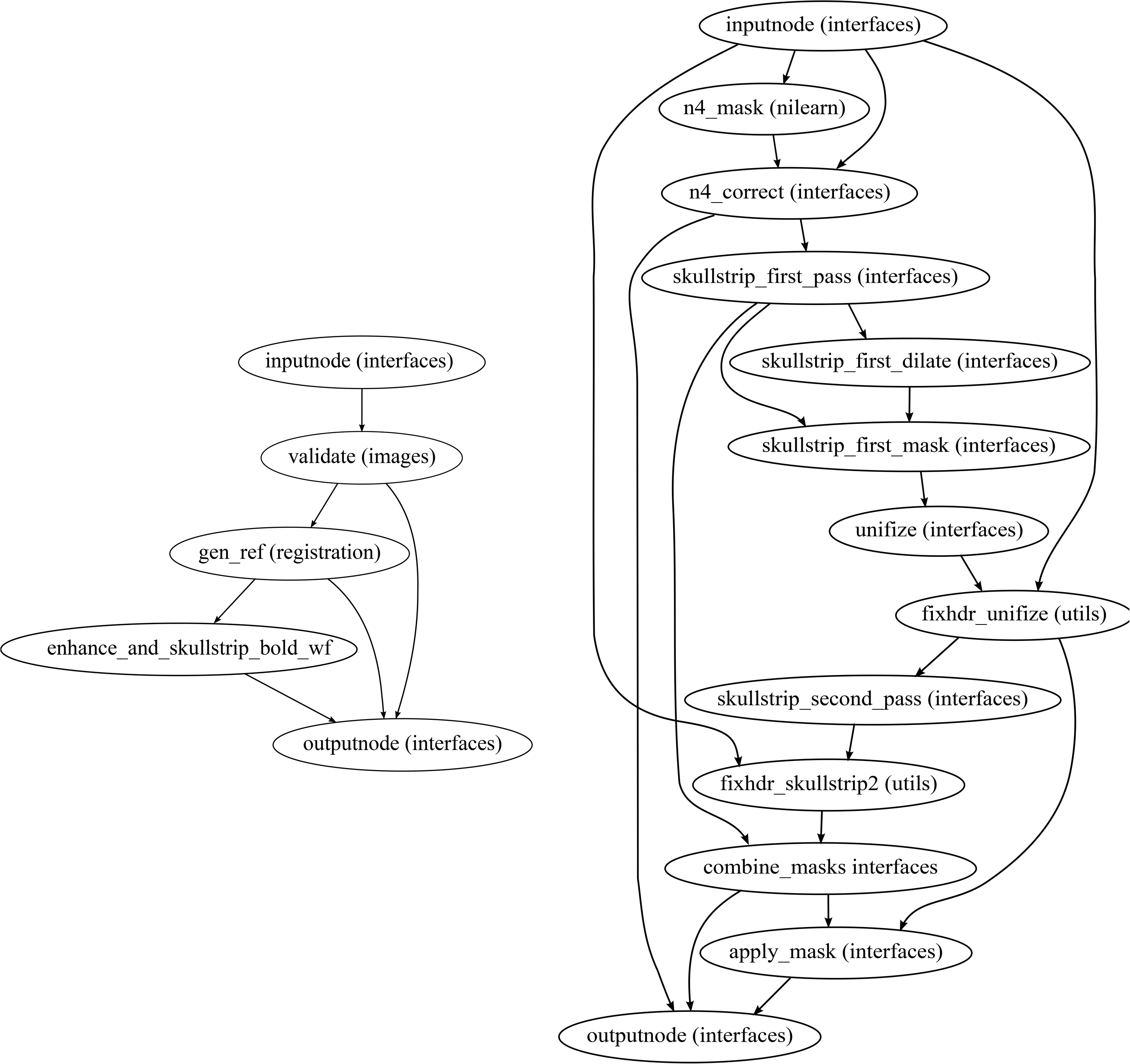

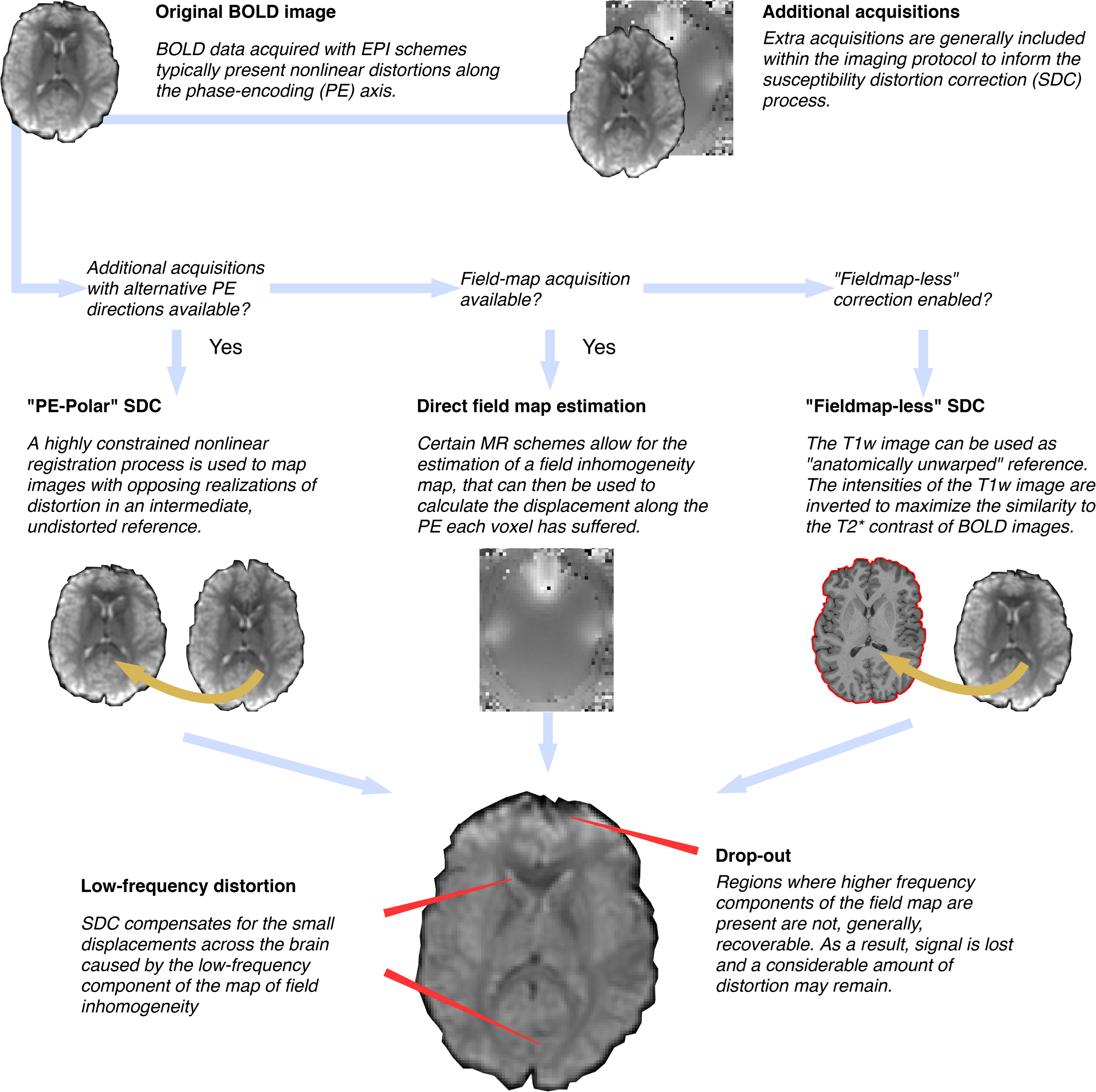

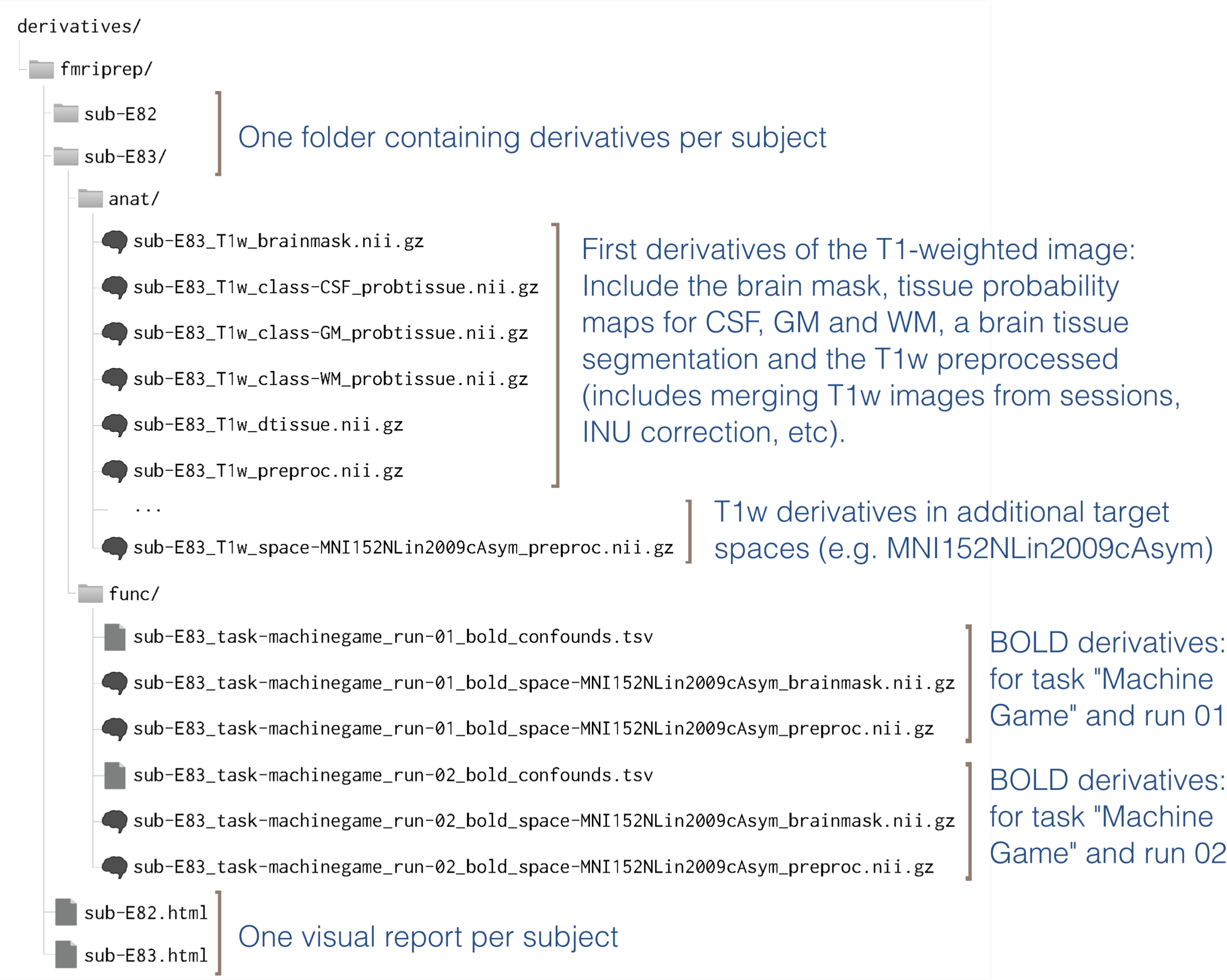

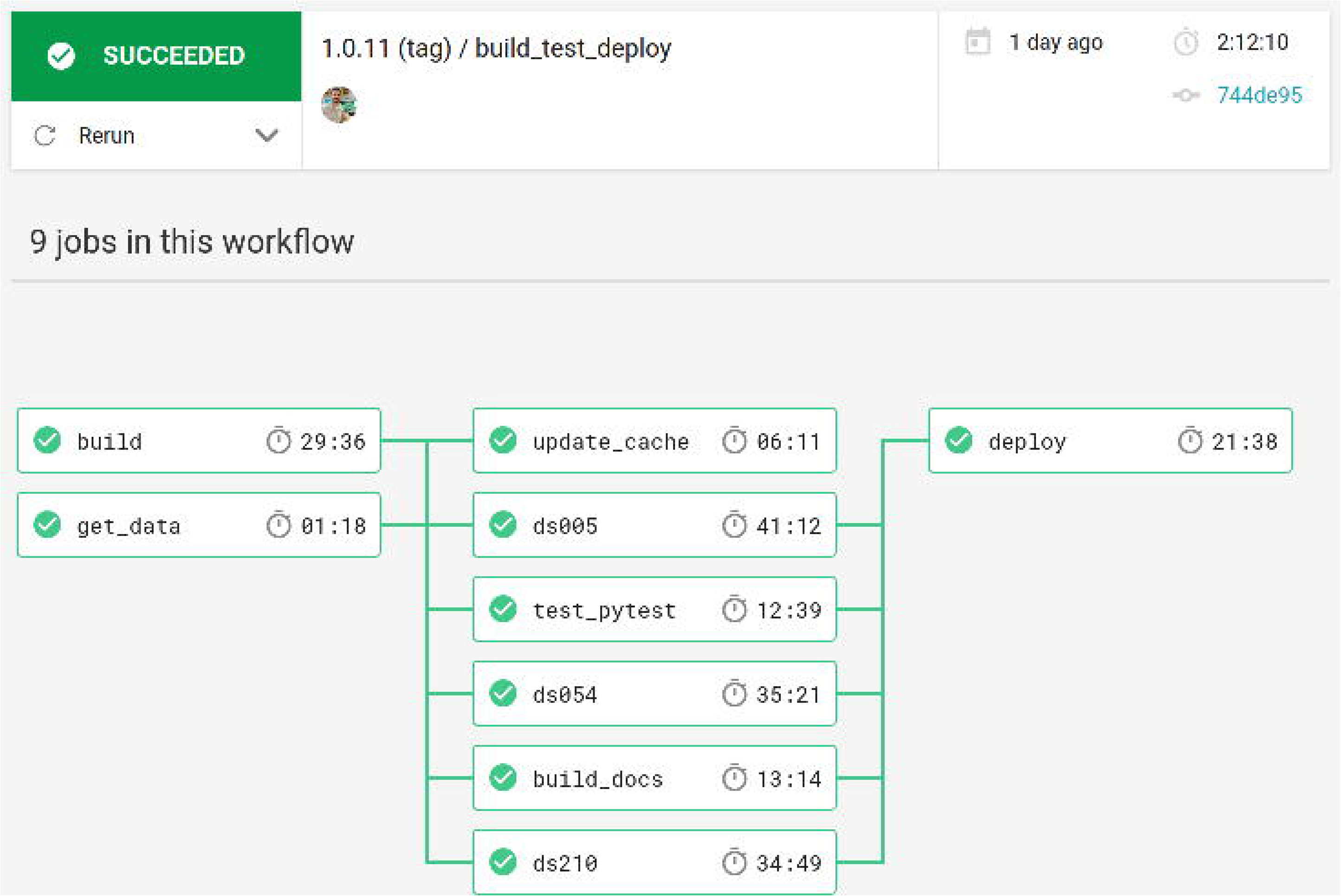

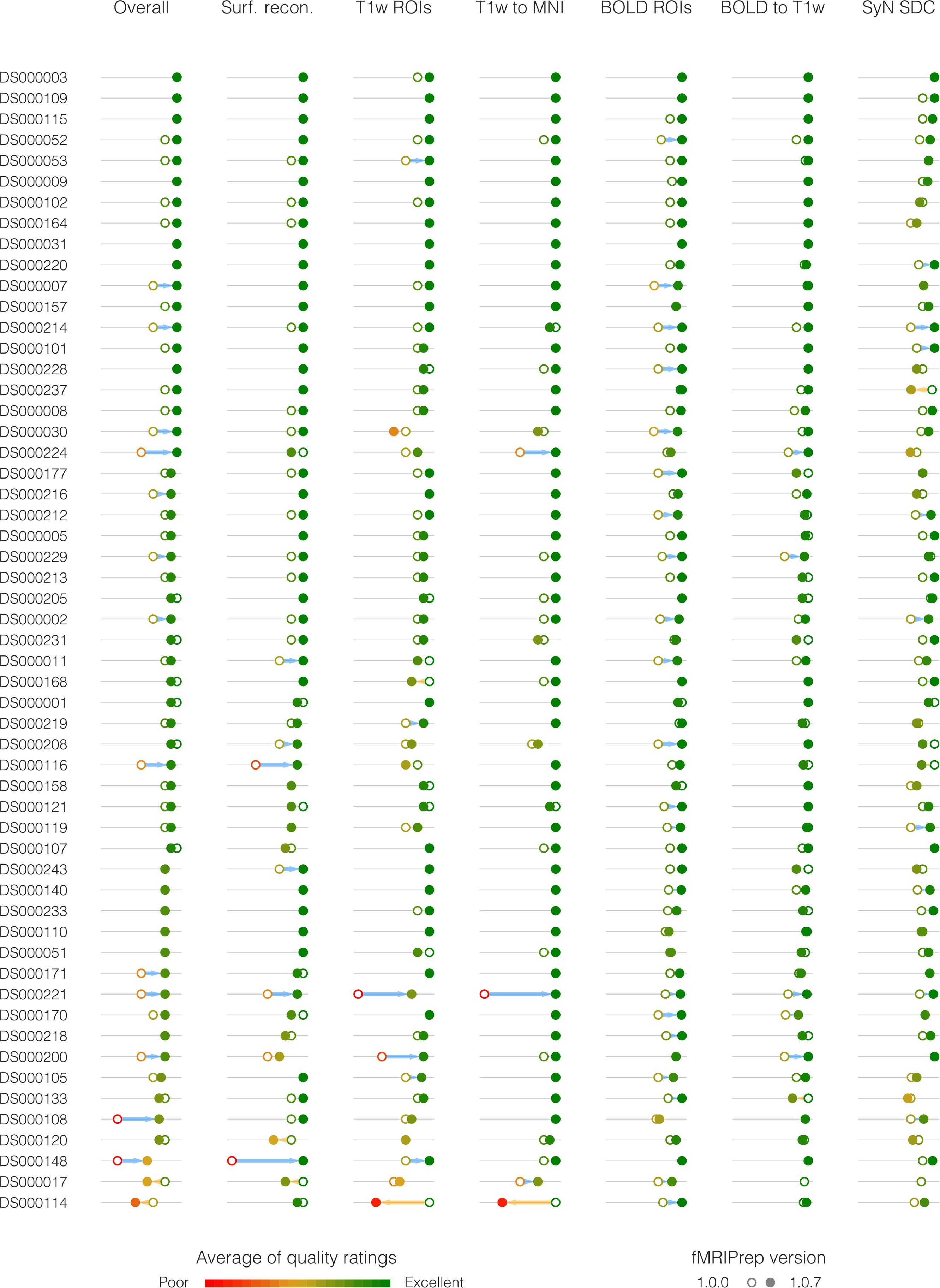

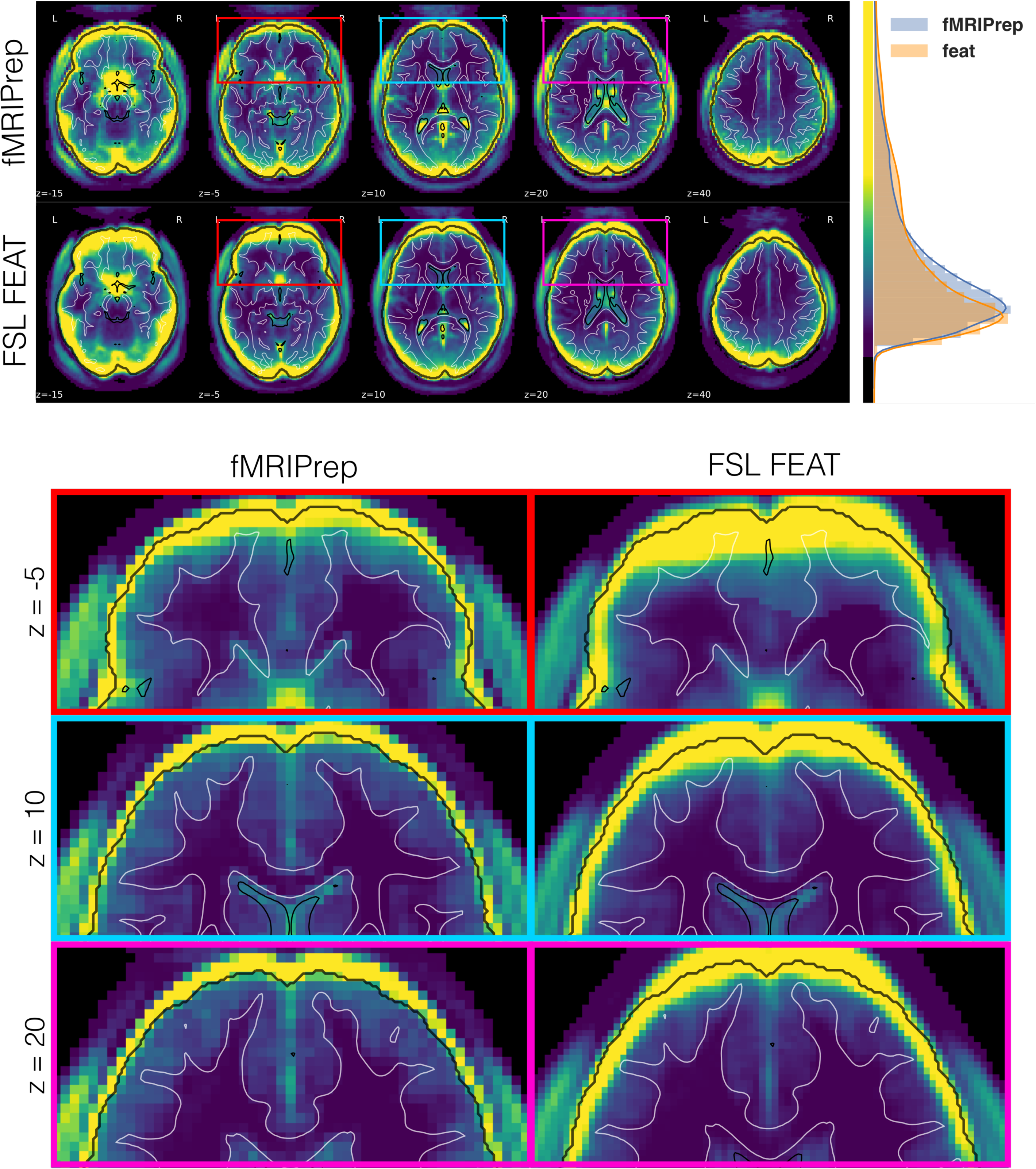

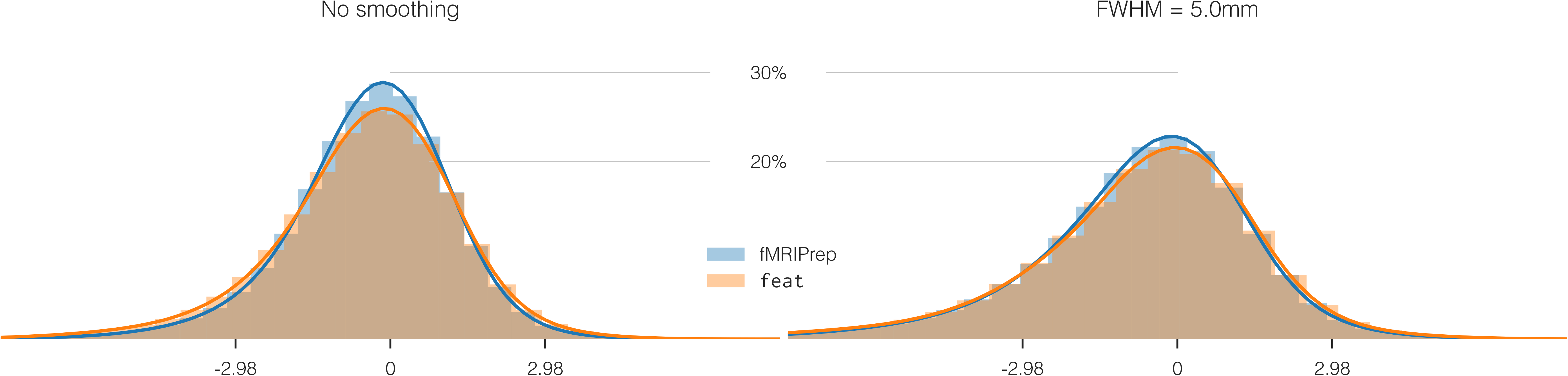

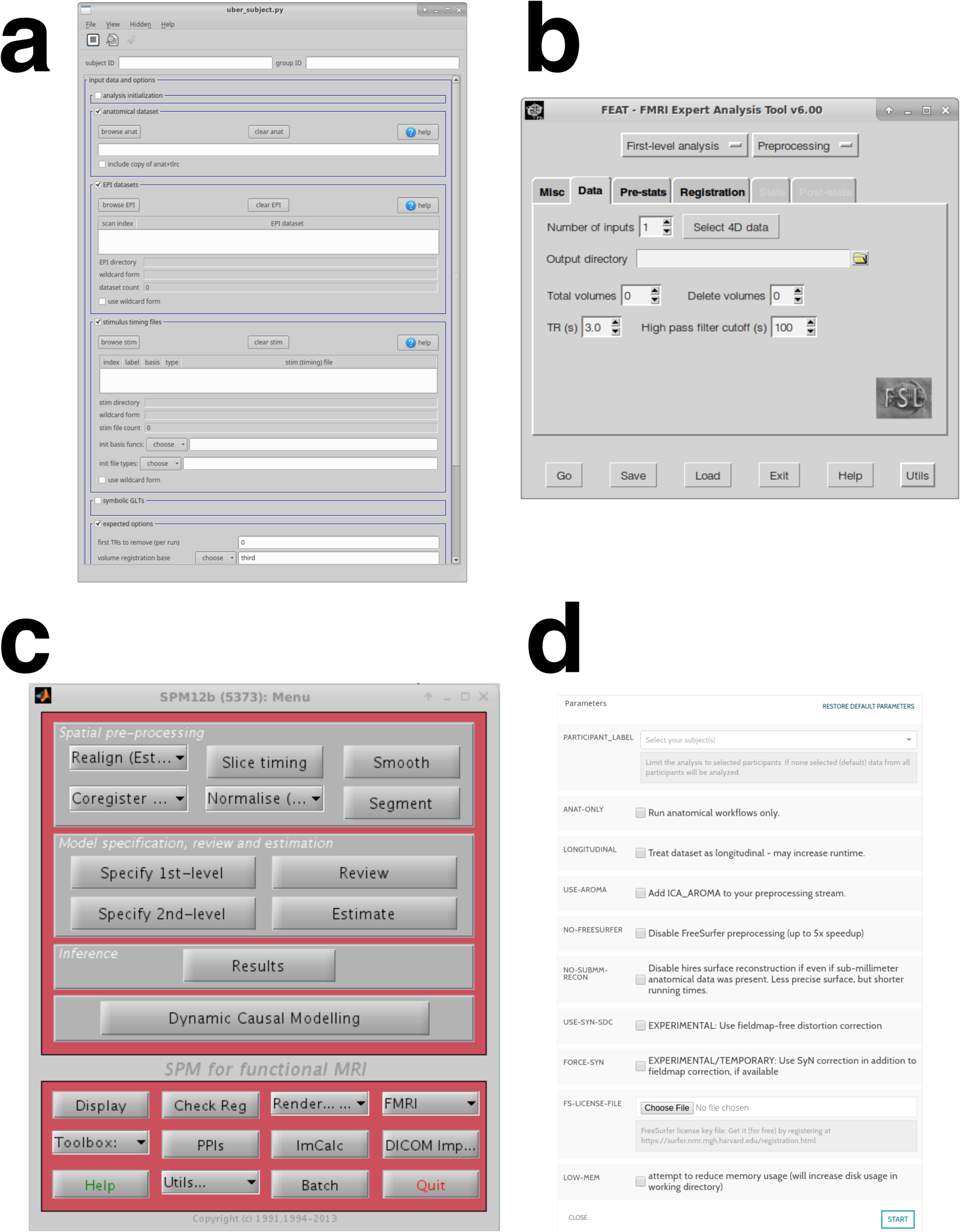

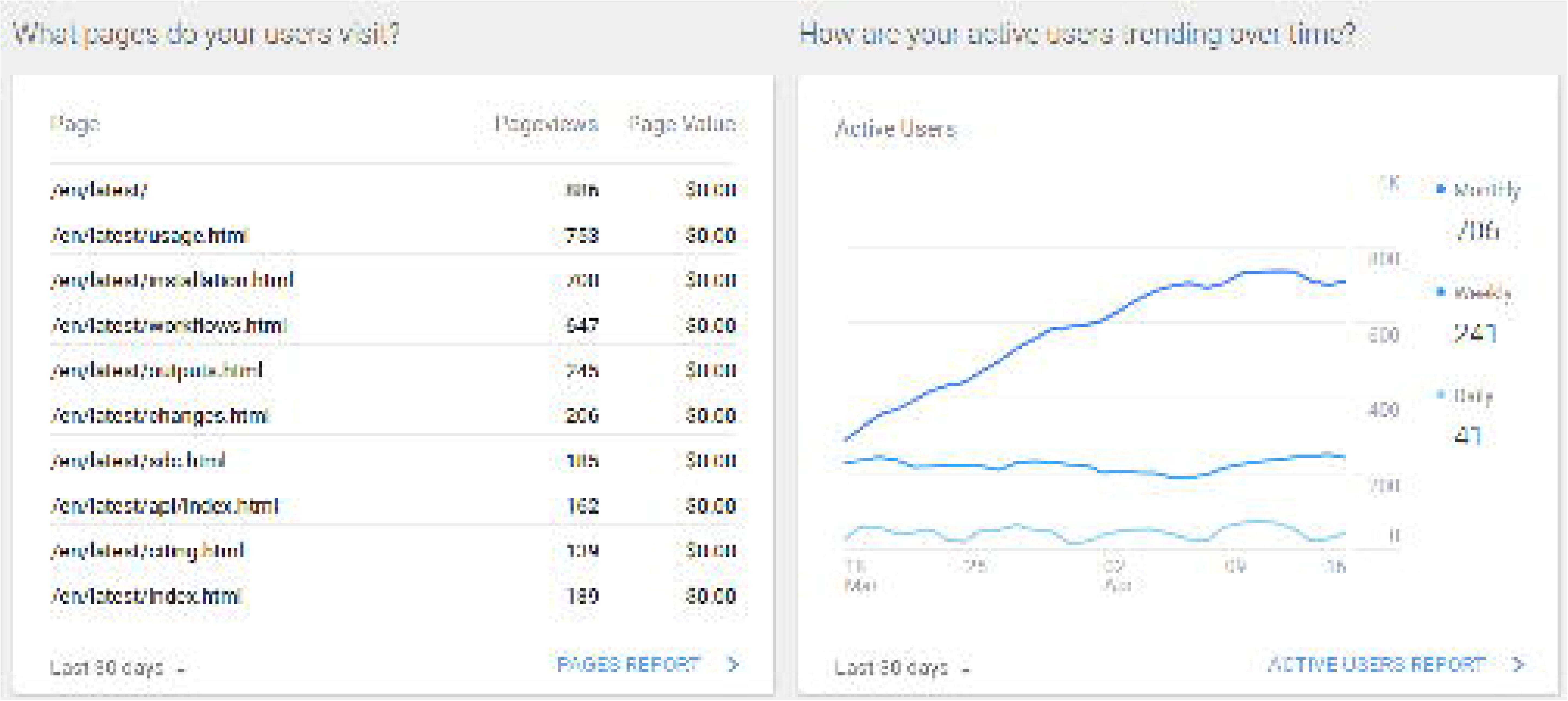

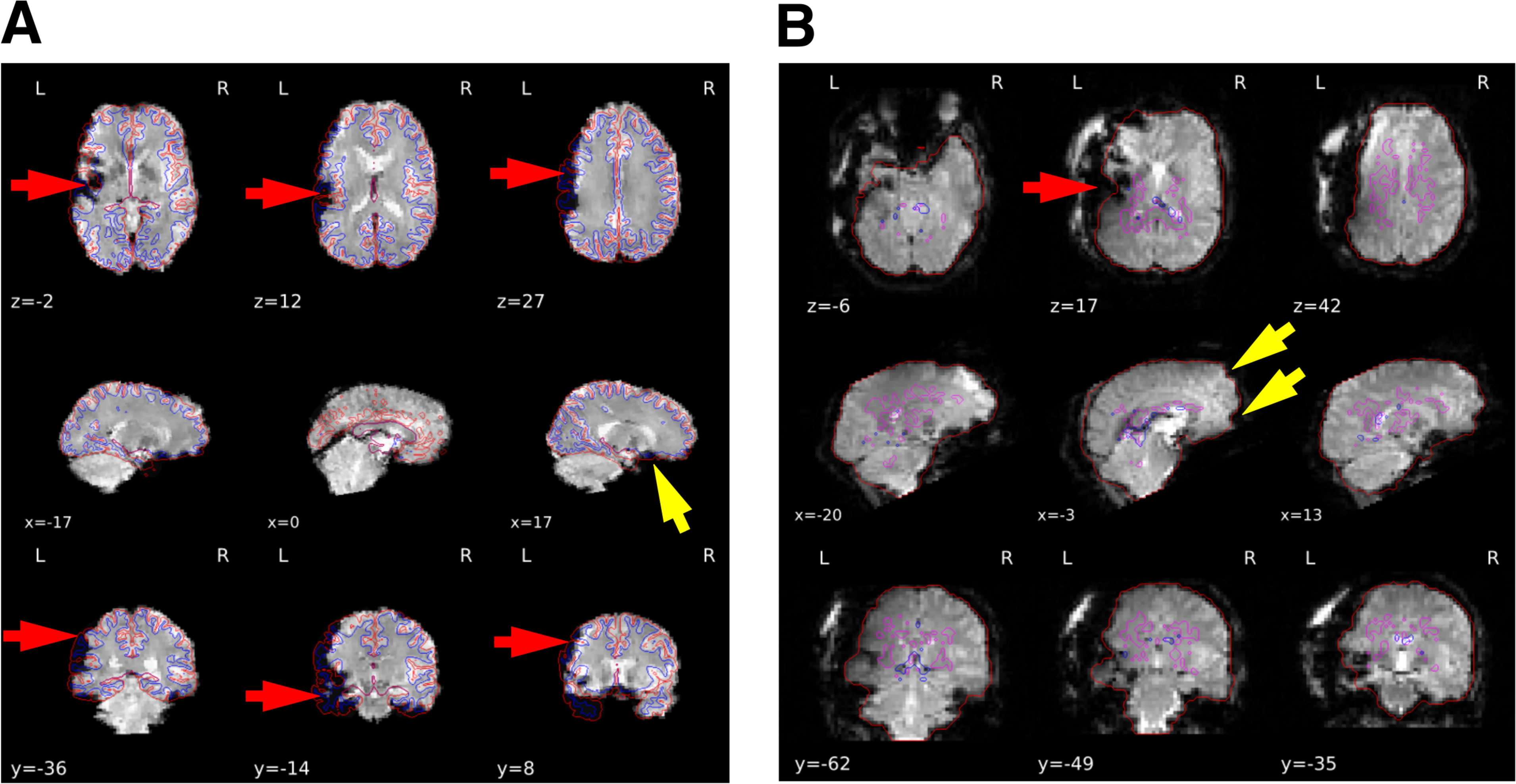

## REFERENCES

1 Poldrack, R. A. & Farah, M. J. Progress and challenges in probing the human brain. Nature 526, 371–379 (2015). doi:10.1038/nature15692.

2 Power, J. D., Plitt, M., Laumann, T. O. & Martin, A. Sources and implications of whole-brain fMRI signals in humans. NeuroImage 146, 609–625 (2017). doi:10.1016/j.neuroimage.2016.09.038.

3 Lindquist, M. A. The Statistical Analysis of fMRI Data. Statistical Science 23, 439–464 (2008). doi:10.1214/09-STS282.

4 Strother, S. C. Evaluating fMRI preprocessing pipelines. IEEE Engineering in Medicine and Biology Magazine 25, 27–41 (2006). doi:10.1109/MEMB.2006.1607667.

5 Sladky, R. et al. Slice-timing effects and their correction in functional MRI. NeuroImage 58, 588–594 (2011). doi:10.1016/j.neuroimage.2011.06.078.

6 Ashburner, J. Preparing fMRI Data for Statistical Analysis. In Filippi, M. (ed.) fMRI Techniques and Protocols, no. 41 in Neuromethods, 151–178 (Humana Press, 2009). doi:10.1007/978-1-60327-919-2_6.

7 Power, J. D., Barnes, K. A., Snyder, A. Z., Schlaggar, B. L. & Petersen, S. E. Spurious but systematic correlations in functional connectivity MRI networks arise from subject motion. NeuroImage 59, 2142–2154 (2012). doi:10.1016/j.neuroimage.2011.10.018.

8 Murphy, K., Birn, R. M., Handwerker, D. A., Jones, T. B. & Bandettini, P. A. The impact of global signal regression on resting state correlations: Are anti-correlated networks introduced? NeuroImage 44, 893–905 (2009). doi:10.1016/j.neuroimage.2008.09.036.

9 Liu, T. T., Nalci, A. & Falahpour, M. The global signal in fMRI: Nuisance or Information? NeuroImage 150, 213–229 (2017). doi:10.1016/j.neuroimage.2017.02.036.

10 Behzadi, Y., Restom, K., Liau, J. & Liu, T. T. A component based noise correction method (CompCor) for BOLD and perfusion based fMRI. NeuroImage 37, 90–101 (2007). doi:10.1016/j.neuroimage.2007.04.042.

11 Pruim, R. H. R. et al. ICA-AROMA: A robust ICA-based strategy for removing motion artifacts from fMRI data. NeuroImage 112, 267–277 (2015). doi:10.1016/j.neuroimage.2015.02.064.

12 Cox, R. W. & Hyde, J. S. Software tools for analysis and visualization of fMRI data. NMR in Biomedicine 10, 171–178 (1997). doi:10.1002/(SICI)1099-1492(199706/08)10:4/5<171::AID-NBM453>3.0.œ;2-L.

13 Avants, B. B. et al. A reproducible evaluation of ANTs similarity metric performance in brain image registration. NeuroImage 54, 2033–44 (2011). doi:10.1016/j.neuroimage.2010.09.025.

14 Fischl, B. FreeSurfer. NeuroImage 62, 774–781 (2012). doi:10.1016/j.neuroimage.2012.01.021.

15 Jenkinson, M., Beckmann, C. F., Behrens, T. E., Woolrich, M. W. & Smith, S. M. FSL. NeuroImage 62, 782–790 (2012). doi:10.1016/j.neuroimage.2011.09.015.

16 Abraham, A. et al. Machine learning for neuroimaging with scikit-learn. Frontiers in Neuroinformatics 8 (2014). doi:10.3389/fninf.2014.00014.

17 Friston, K. J., Ashburner, J., Kiebel, S. J., Nichols, T. E. & Penny, W. D. Statistical parametric mapping: the analysis of functional brain images (Academic Press, London, 2006).

18 Power, J. D., Plitt, M., Kundu, P., Bandettini, P. A. & Martin, A. Temporal interpolation alters motion in fMRI scans: Magnitudes and consequences for artifact detection. PLOS ONE 12, e0182939 (2017). doi:10.1371/journal.pone.0182939.

19 Carp, J. The secret lives of experiments: Methods reporting in the fMRI literature. NeuroImage 63, 289–300 (2012). doi:10.1016/j.neuroimage.2012.07.004.

20 Van Essen, D. et al. The Human Connectome Project: A data acquisition perspective. NeuroImage 62, 2222–2231 (2012). doi:10.1016/j.neuroimage.2012.02.018.

21 Miller, K. L. et al. Multimodal population brain imaging in the UK Biobank prospective epidemiological study. Nature Neuroscience 19, 1523–1536 (2016). doi:10.1038/nn.4393.

22 Glasser, M. F. et al. The minimal preprocessing pipelines for the Human Connectome Project. NeuroImage 80, 105–124 (2013). doi:10.1016/j.neuroimage.2013.04.127.

23 Alfaro-Almagro, F. et al. Image processing and Quality Control for the first 10,000 brain imaging datasets from UK Biobank. NeuroImage (2017). doi:10.1016/j.neuroimage.2017.10.034.

24 Gorgolewski, K. J. et al. The brain imaging data structure, a format for organizing and describing outputs of neuroimaging experiments. Scientific Data 3, 160044 (2016). doi:10.1038/sdata.2016.44.

25 Gorgolewski, K. J. et al. Nipype: a flexible, lightweight and extensible neuroimaging data processing framework in Python. Zenodo [Software] (2016). doi:10.5281/zenodo.50186.

26 Tustison, N. J. et al. N4ITK: Improved N3 Bias Correction. IEEE Transactions on Medical Imaging 29, 1310–1320 (2010). doi:10.1109/TMI.2010.2046908.

27 Marcus, D. S. et al. Open Access Series of Imaging Studies (OASIS): Cross-sectional MRI Data in Young, Middle Aged, Nondemented, and Demented Older Adults. Journal of Cognitive Neuroscience 19, 1498–1507 (2007). doi:10.1162/jocn.2007.19.9.1498.

28 Nooner, K. B. et al. The NKI-Rockland Sample: A Model for Accelerating the Pace of Discovery Science in Psychiatry. Frontiers in Neuroscience 6 (2012). doi:10.3389/fnins.2012.00152.

29 Reuter, M., Rosas, H. D. & Fischl, B. Highly accurate inverse consistent registration: A robust approach. NeuroImage 53, 1181–1196 (2010). doi:10.1016/j.neuroimage.2010.07.020.

30 Dale, A. M., Fischl, B. & Sereno, M. I. Cortical Surface-Based Analysis: I. Segmentation and Surface Reconstruction. NeuroImage 9, 179–194 (1999). doi:10.1006/nimg.1998.0395.

31 Klein, A. et al. Mindboggling morphometry of human brains. PLOS Computational Biology 13, e1005350 (2017). doi:10.1371/journal.pcbi.1005350.

32 Fonov, V., Evans, A., McKinstry, R., Almli, C. & Collins, D. Unbiased nonlinear average age-appropriate brain templates from birth to adulthood. NeuroImage 47, Supplement 1, S102 (2009). doi:10.1016/S1053-8119(09)70884-5.

33 Avants, B., Epstein, C., Grossman, M. & Gee, J. Symmetric diffeomorphic image registration with cross-correlation: Evaluating automated labeling of elderly and neurodegenerative brain. Medical Image Analysis 12, 26–41 (2008). doi:10.1016/j.media.2007.06.004.

34 Klein, A. et al. Evaluation of 14 nonlinear deformation algorithms applied to human brain MRI registration. NeuroImage 46, 786–802 (2009). doi:doi:10.1016/j.neuroimage.2008.12.037.

35 Zhang, Y., Brady, M. & Smith, S. Segmentation of brain MR images through a hidden Markov random field model and the expectation-maximization algorithm. IEEE Transactions on Medical Imaging 20, 45–57 (2001). doi:10.1109/42.906424.

36 Jenkinson, M., Bannister, P., Brady, M. & Smith, S. Improved Optimization for the Robust and Accurate Linear Registration and Motion Correction of Brain Images. NeuroImage 17, 825–841 (2002). doi:10.1006/nimg.2002.1132.

37 Oakes, T. R. et al. Comparison of fMRI motion correction software tools. NeuroImage 28, 529–543 (2005). doi:10.1016/j.neuroimage.2005.05.058.

38 Greve, D. N. & Fischl, B. Accurate and robust brain image alignment using boundary-based registration. NeuroImage 48, 63–72 (2009). doi:10.1016/j.neuroimage.2009.06.060.

39 Lanczos, C. Evaluation of Noisy Data. Journal of the Society for Industrial and Applied Mathematics Series B Numerical Analysis 1, 76–85 (1964). doi:10.1137/0701007.

40 Power, J. D. et al. Methods to detect, characterize, and remove motion artifact in resting state fMRI. NeuroImage 84, 320–341 (2014). doi:10.1016/j.neuroimage.2013.08.048.

41 Poldrack, R. A. et al. Guidelines for reporting an fMRI study. NeuroImage 40, 409–414 (2008). doi:10.1016/j.neuroimage.2007.11.048.

42 Sikka, S. et al. Towards automated analysis of connectomes: The configurable pipeline for the analysis of connectomes (C-PAC). In 5th INCF Congress of Neuroinformatics, vol. 117 (Munich, Germany, 2014). doi:10.3389/conf.fninf.2014.08.00117.

43 Wang, S. et al. Evaluation of Field Map and Nonlinear Registration Methods for Correction of Susceptibility Artifacts in Diffusion MRI. Frontiers in Neuroinformatics 11 (2017). doi:10.3389/fninf.2017.00017.

44 McIntosh, S., Kamei, Y., Adams, B. & Hassan, A. E. The Impact of Code Review Coverage and Code Review Participation on Software Quality: A Case Study of the Qt, VTK, and ITK Projects. In Proceedings of the 11th Working Conference on Mining Software Repositories, MSR 2014, 192–201 (ACM, New York, NY, USA, 2014). doi:10.1145/2597073.2597076.

45 Gorgolewski, K. J. et al. BIDS Apps: Improving ease of use, accessibility, and reproducibility of neuroimaging data analysis methods. PLOS Computational Biology 13, e1005209 (2017). doi:10.1371/journal.pcbi.1005209.

46 Beaulieu-Jones, B. K. & Greene, C. S. Reproducibility of computational workflows is automated using continuous analysis. Nature Biotechnology 35, 342 (2017). doi:10.1038/nbt.3780.

47 Kurtzer, G. M., Sochat, V. & Bauer, M. W. Singularity: Scientific containers for mobility of compute. PLOS ONE 12, e0177459 (2017). doi:10.1371/journal.pone.0177459.

48 Schonberg, T. et al. Decreasing Ventromedial Prefrontal Cortex Activity During Sequential Risk-Taking: An fMRI Investigation of the Balloon Analog Risk Task. Frontiers in Neuroscience 6 (2012). doi:10.3389/fnins.2012.00080.

49 Aron, A. R., Gluck, M. A. & Poldrack, R. A. Long-term test-retest reliability of functional MRI in a classification learning task. NeuroImage 29, 1000–1006 (2006). doi:10.1016/j.neuroimage.2005.08.010.

50 Xue, G. & Poldrack, R. A. The Neural Substrates of Visual Perceptual Learning of Words: Implications for the Visual Word Form Area Hypothesis. Journal of Cognitive Neuroscience 19, 1643–1655 (2007). doi:10.1162/jocn.2007.19.10.1643.

51 Tom, S. M., Fox, C. R., Trepel, C. & Poldrack, R. A. The Neural Basis of Loss Aversion in Decision-Making Under Risk. Science 315,515–518 (2007). doi:10.1126/science.1134239.

52 Xue, G., Aron, A. R. & Poldrack, R. A. Common Neural Substrates for Inhibition of Spoken and Manual Responses. Cerebral Cortex 18, 1923–1932 (2008). doi:10.1093/cercor/bhm220.

53 Aron, A. R., Behrens, T. E., Smith, S., Frank, M. J. & Poldrack, R. A. Triangulating a Cognitive Control Network Using Diffusion-Weighted Magnetic Resonance Imaging (MRI) and Functional MRI. Journal of Neuroscience 27, 3743–3752 (2007). doi:10.1523/JNEUROSCI.0519-07.2007.

54 Foerde, K., Knowlton, B. J. & Poldrack, R. A. Modulation of competing memory systems by distraction. Proceedings of the National Academy of Sciences 103, 11778–11783 (2006). doi:10.1073/pnas.0602659103.

55 Poldrack, R. A. et al. A phenome-wide examination of neural and cognitive function. Scientific Data 3, 160110 (2016). doi:10.1038/sdata.2016.110.

56 Gorgolewski, K. J., Durnez, J. & Poldrack, R. A. Preprocessed Consortium for Neuropsychiatric Phenomics dataset. F1000Research 6, 1262 (2017). doi:10.12688/f1000research.11964.2.

57 Laumann, T. O. et al. Functional System and Areal Organization of a Highly Sampled Individual Human Brain. Neuron 87, 657–670 (2015). doi:10.1016/j.neuron.2015.06.037.

58 Alvarez, R., Jasdzewski, G. & Poldrack, R. A. Building memories in two languages: an fMRI study of episodic encoding in bilinguals. In SfN Neuroscience (Orlando, FL, US, 2002). URL http://www.sfn.org/annual-meeting/past-and-future-annual-meetings/abstract-archive/abstract-archive-detail.

59 Poldrack, R. A. et al. Interactive memory systems in the human brain. Nature 414, 546–550 (2001). doi:10.1038/35107080.

60 Kelly, A. M. C., Uddin, L. Q., Biswal, B. B., Castellanos, F. X. & Milham, M. P. Competition between functional brain networks mediates behavioral variability. NeuroImage 39, 527–537 (2008). doi:10.1016/j.neuroimage.2007.08.008.

61 Mennes, M. et al. Inter-individual differences in resting-state functional connectivity predict task-induced BOLD activity. NeuroImage 50, 1690–1701 (2010). doi:10.1016/j.neuroimage.2010.01.002.

62 Mennes, M. et al. Linking inter-individual differences in neural activation and behavior to intrinsic brain dynamics. NeuroImage 54, 2950–2959 (2011). doi:10.1016/j.neuroimage.2010.10.046.

63 Haxby, J. V. et al. Distributed and Overlapping Representations of Faces and Objects in Ventral Temporal Cortex. Science 293, 2425–2430 (2001). doi:10.1126/science.1063736.

64 Hanson, S. J., Matsuka, T. & Haxby, J. V. Combinatorial codes in ventral temporal lobe for object recognition: Haxby (2001) revisited: is there a âĂIJfaceâAi area? NeuroImage 23, 156–166 (2004). doi:10.1016/j.neuroimage.2004.05.020.

65 Duncan, K. J., Pattamadilok, C., Knierim, I. & Devlin, J. T. Consistency and variability in functional localisers. NeuroImage 46, 1018–1026 (2009). doi:10.1016/j.neuroimage.2009.03.014.

66 Wager, T. D., Davidson, M. L., Hughes, B. L., Lindquist, M. A. & Ochsner, K. N. Prefrontal-Subcortical Pathways Mediating Successful Emotion Regulation. Neuron 59, 1037–1050 (2008). doi:10.1016/j.neuron.2008.09.006.

67 Moran, J. M., Jolly, E. & Mitchell, J. P. Social-Cognitive Deficits in Normal Aging. Journal of Neuroscience 32, 5553–5561 (2012). doi:10.1523/JNEUROSCI.5511-11.2012.

68 Uncapher, M. R., Hutchinson, J. B. & Wagner, A. D. Dissociable Effects of Top-Down and Bottom-Up Attention during Episodic Encoding. Journal of Neuroscience 31, 12613–12628 (2011). doi:10.1523/JNEUROSCI.0152-11.2011.

69 Gorgolewski, K. J. et al. A test-retest fMRI dataset for motor, language and spatial attention functions. GigaScience 2, 1–4 (2013). doi:10.1186/2047-217X-2-6.

70 Repovs, G. & Barch, D. M. Working memory related brain network connectivity in individuals with schizophrenia and their siblings. Frontiers in Human Neuroscience 6 (2012). doi:10.3389/fnhum.2012.00137.

71 Repovs, G., Csernansky, J. G. & Barch, D. M. Brain Network Connectivity in Individuals with Schizophrenia and Their Siblings. Biological Psychiatry 69, 967–973 (2011). doi:10.1016/j.biopsych.2010.11.009.

72 Walz, J. M. et al. Simultaneous EEG-fMRI Reveals Temporal Evolution of Coupling between Supramodal Cortical Attention Networks and the Brainstem. Journal of Neuroscience 33, 19212–19222 (2013). doi:10.1523/JNEUROSCI.2649-13.2013.

73 Walz, J. M. et al. Simultaneous EEG-fMRI reveals a temporal cascade of task-related and default-mode activations during a simple target detection task. NeuroImage 102, 229–239 (2014). doi:10.1016/j.neuroimage.2013.08.014.

74 Conroy, B. R., Walz, J. M. & Sajda, P. Fast Bootstrapping and Permutation Testing for Assessing Reproducibility and Interpretability of Multivariate fMRI Decoding Models. PLOS ONE 8, e79271 (2013). doi:10.1371/journal.pone.0079271.

75 Walz, J. M. et al. Prestimulus EEG alpha oscillations modulate task-related fMRI BOLD responses to auditory stimuli. NeuroImage 113, 153–163 (2015). doi:10.1016/j.neuroimage.2015.03.028.

76 Velanova, K., Wheeler, M. E. & Luna, B. Maturational Changes in Anterior Cingulate and Frontoparietal Recruitment Support the Development of Error Processing and Inhibitory Control. Cerebral Cortex 18, 2505–2522 (2008). doi:10.1093/cercor/bhn012.

77 Padmanabhan, A., Geier, C. F., Ordaz, S. J., Teslovich, T. & Luna, B. Developmental changes in brain function underlying the influence of reward processing on inhibitory control. Developmental Cognitive Neuroscience 1, 517–529 (2011). doi:10.1016/j.dcn.2011.06.004.

78 Geier, C. F., Terwilliger, R., Teslovich, T., Velanova, K. & Luna, B. Immaturities in Reward Processing and Its Influence on Inhibitory Control in Adolescence. Cerebral Cortex 20, 1613–1629 (2010). doi:10.1093/cercor/bhp225.

79 Cera, N., Tartaro, A. & Sensi, S. L. Modafinil Alters Intrinsic Functional Connectivity of the Right Posterior Insula: A Pharmacological Resting State fMRI Study. PLOS ONE 9, e107145 (2014). doi:10.1371/journal.pone.0107145.

80 Woo, C.-W., Roy, M., Buhle, J. T. & Wager, T. D. Distinct Brain Systems Mediate the Effects of Nociceptive Input and Self-Regulation on Pain. PLOS Biology 13, e1002036 (2015). doi:10.1371/journal.pbio.1002036.

81 Smeets, P. A. M., Kroese, F. M., Evers, C. & de Ridder, D. T. D. Allured or alarmed: Counteractive control responses to food temptations in the brain. Behavioural Brain Research 248, 41–45 (2013). doi:10.1016/j.bbr.2013.03.041.

82 Pernet, C. R. et al. The human voice areas: Spatial organization and inter-individual variability in temporal and extra-temporal cortices. NeuroImage 119, 164–174 (2015). doi:10.1016/j.neuroimage.2015.06.050.

83 Verstynen, T. D. The organization and dynamics of corticostriatal pathways link the medial orbitofrontal cortex to future behavioral responses. Journal of Neurophysiology 112, 2457–2469 (2014). doi:10.1152/jn.00221.2014.

84 Bursley, J. K., Nestor, A., Tarr, M. J. & Creswell, J. D. Awake, Offline Processing during Associative Learning. PLOS ONE 11, e0127522 (2016). doi:10.1371/journal.pone.0127522.

85 Ella, G., David, M. & Avi, K. Learning from the other limb’s experience: sharing the âAŸtrainedâĂZ M1 representation of the motor sequence knowledge. The Journal of Physiology 594, 169–188 (2015). doi:10.1113/JP270184.

86 Gabitov, E., Manor, D. & Karni, A. Patterns of Modulation in the Activity and Connectivity of Motor Cortex during the Repeated Generation of Movement Sequences. Journal of Cognitive Neuroscience 27, 736–751 (2014). doi:10.1162/jocn_a_00751.

87 Gabitov, E., Manor, D. & Karni, A. Done That: Short-term Repetition Related Modulations of Motor Cortex Activity as a Stable Signature for Overnight Motor Memory Consolidation. Journal of Cognitive Neuroscience 26, 2716–2734 (2014). doi:10.1162/jocn_a_00675.

88 Lepping, R. J., Atchley, R. A. & Savage, C. R. Development of a validated emotionally provocative musical stimulus set for research. Psychology of Music 44, 1012–1028 (2016). doi:10.1177/0305735615604509.

89 Park, C.-A. & Kang, C.-K. Sensing the effects of mouth breathing by using 3-tesla MRI. Journal of the Korean Physical Society 70, 1070–1076 (2017). doi:10.3938/jkps.70.1070.

90 Iannilli, E. et al. Effects of Manganese Exposure on Olfactory Functions in Teenagers: A Pilot Study. PLOS ONE 11, e0144783 (2016). doi:10.1371/journal.pone.0144783.

91 Kim, J., Wang, J., Wedell, D. H. & Shinkareva, S. V. Identifying Core Affect in Individuals from fMRI Responses to Dynamic Naturalistic Audiovisual Stimuli. PLOS ONE 11, e0161589 (2016). doi:10.1371/journal.pone.0161589.

92 TÃl’treault, P. et al. Brain Connectivity Predicts Placebo Response across Chronic Pain Clinical Trials. PLOS Biology 14, e1002570 (2016). doi:10.1371/journal.pbio.1002570.

93 Chakroff, A. et al. When minds matter for moral judgment: intent information is neurally encoded for harmful but not impure acts. Social Cognitive and Affective Neuroscience 11, 476–484 (2016). doi:10.1093/scan/nsv131.

94 Koster-Hale, J., Saxe, R., Dungan, J. & Young, L. L. Decoding moral judgments from neural representations of intentions. Proceedings of the National Academy of Sciences 110, 5648–5653 (2013). doi:10.1073/pnas.1207992110.

95 Gao, X. et al. My Body Looks Like That GirlâAZs: Body Mass Index Modulates Brain Activity during Body Image Self-Reflection among Young Women. PLOS ONE 11, e0164450 (2016). doi:10.1371/journal.pone.0164450.

96 Romaniuk, L., Pope, M., Nicol, K., Steele, D. & Hall, J. Neural correlates of fears of abandonment and rejection in borderline personality disorder. Wellcome Open Research 1, 33 (2016). doi:10.12688/wellcomeopenres.10331.1.

97 Cohen, A. D., Nencka, A. S., Lebel, R. M. & Wang, Y. Multiband multi-echo imaging of simultaneous oxygenation and flow timeseries for resting state connectivity. PLOS ONE 12, e0169253 (2017). doi:10.1371/journal.pone.0169253.

98 Dalenberg, J. R., Weitkamp, L., Renken, R. J., Nanetti, L. & Horst, G. J. t. Flavor pleasantness processing in the ventral emotion network. PLOS ONE 12, e0170310 (2017). doi:10.1371/journal.pone.0170310.

99 Roy, A. et al. The evolution of cost-efficiency in neural networks during recovery from traumatic brain injury. PLOS ONE 12, e0170541 (2017). doi:10.1371/journal.pone.0170541.

100 Gordon, E. M. et al. Precision Functional Mapping of Individual Human Brains. Neuron 95, 791–807.e7 (2017). doi:10.1016/j.neuron.2017.07.011.

101 Veldhuizen, M. G. et al. Integration of Sweet Taste and Metabolism Determines Carbohydrate Reward. Current Biology 27, 2476–2485.e6 (2017). doi:10.1016/j.cub.2017.07.018.

102 Greene, D. J. et al. Behavioral interventions for reducing head motion during MRI scans in children. NeuroImage 171, 234–245 (2018). doi:10.1016/j.neuroimage.2018.01.023.

103 Nastase, S. A. et al. Attention Selectively Reshapes the Geometry of Distributed Semantic Representation. Cerebral Cortex 27, 4277–4291 (2017). doi:10.1093/cercor/bhx138.

104 Kanazawa, Y. et al. Phonological memory in sign language relies on the visuomotor neural system outside the left hemisphere language network. PLOS ONE 12, e0177599 (2017). doi:10.1371/journal.pone.0177599.

105 Esteban, O. et al. MRIQC: Advancing the automatic prediction of image quality in MRI from unseen sites. PLOS ONE 12, e0184661 (2017). doi:10.1371/journal.pone.0184661.

106 Calhoun, V. D. et al. The impact of T1 versus EPI spatial normalization templates for fMRI data analyses. Human BrainMapping 38, 5331–5342 (2017). doi:10.1002/hbm.23737.

107 Beckmann, C. F., Jenkinson, M. & Smith, S. M. General multilevel linear modeling for group analysis in FMRI. NeuroImage 20, 1052–1063 (2003). doi:10.1016/S1053-8119(03)00435-X.

108 Strother, S. C. et al. The Quantitative Evaluation of Functional Neuroimaging Experiments: The NPAIRS Data Analysis Framework. NeuroImage 15, 747–771 (2002). doi:10.1006/nimg.2001.1034.

109 Karaman, M., Nencka, A. S., Bruce, I. P. & Rowe, D. B. Quantification of the Statistical Effects of Spatiotemporal Processing of Nontask fMRI Data. Brain Connectivity 4, 649–661 (2014). doi:10.1089/brain.2014.0278.

110 Gorgolewski, K. et al. Nipype: a flexible, lightweight and extensible neuroimaging data processing framework in Python. Frontiers in Neuroinformatics 5, 13 (2011). doi:10.3389/fninf.2011.00013.

